# The Alzheimer’s disease risk factor APOE4 drives pro-inflammation in human astrocytes *via* HDAC-dependent repression of TAGLN3

**DOI:** 10.1101/2021.04.16.440108

**Authors:** Laurie Arnaud, Philippe Benech, Louise Greetham, Delphine Stephan, Angélique Jimenez, Nicolas Jullien, Laura García-González, Philipp O. Tsvetkov, François Devred, Ignacio Sancho-Martinez, Juan-Carlos Izpisua Belmonte, Kevin Baranger, Santiago Rivera, Emmanuel Nivet

## Abstract

The Apolipoprotein E4 (*APOE4*) is the major allelic risk factor for late-onset Alzheimer’s disease (AD). APOE4 associates with a pro-inflammatory phenotype increasingly considered as critical in AD initiation and progression. Yet, the mechanisms driving an APOE4-dependent neuroinflammation remain unelucidated. Leveraging patient specific human induced Pluripotent Stem Cells (iPSCs) we demonstrate inflammatory chronicity and hyperactivated responses upon cytokines in human APOE4 astrocytes *via* a novel mechanism. We uncovered that APOE4 represses Transgelin 3 (TAGLN3), a new interacting partner of IκBα, thus increasing the NF-kB activity. The transcriptional repression of TAGLN3 was shown to result from an APOE4-dependent histone deacetylase (HDAC) activity. The functional relevance of TAGLN3 was demonstrated by the attenuation of APOE4-driven neuroinflammation after TAGLN3 supplementation. Importantly, TAGLN3 downregulation was confirmed in the brain of AD patients. Our findings highlight the APOE4-TAGLN3 axis as a new pathogenic pathway that paves the way for the development of therapeutics to prevent maladaptive inflammatory responses in *APOE4* carriers, while placing TAGLN3 downregulation as a potential biomarker of AD.

**GRAPHICAL ABSTRACT:** 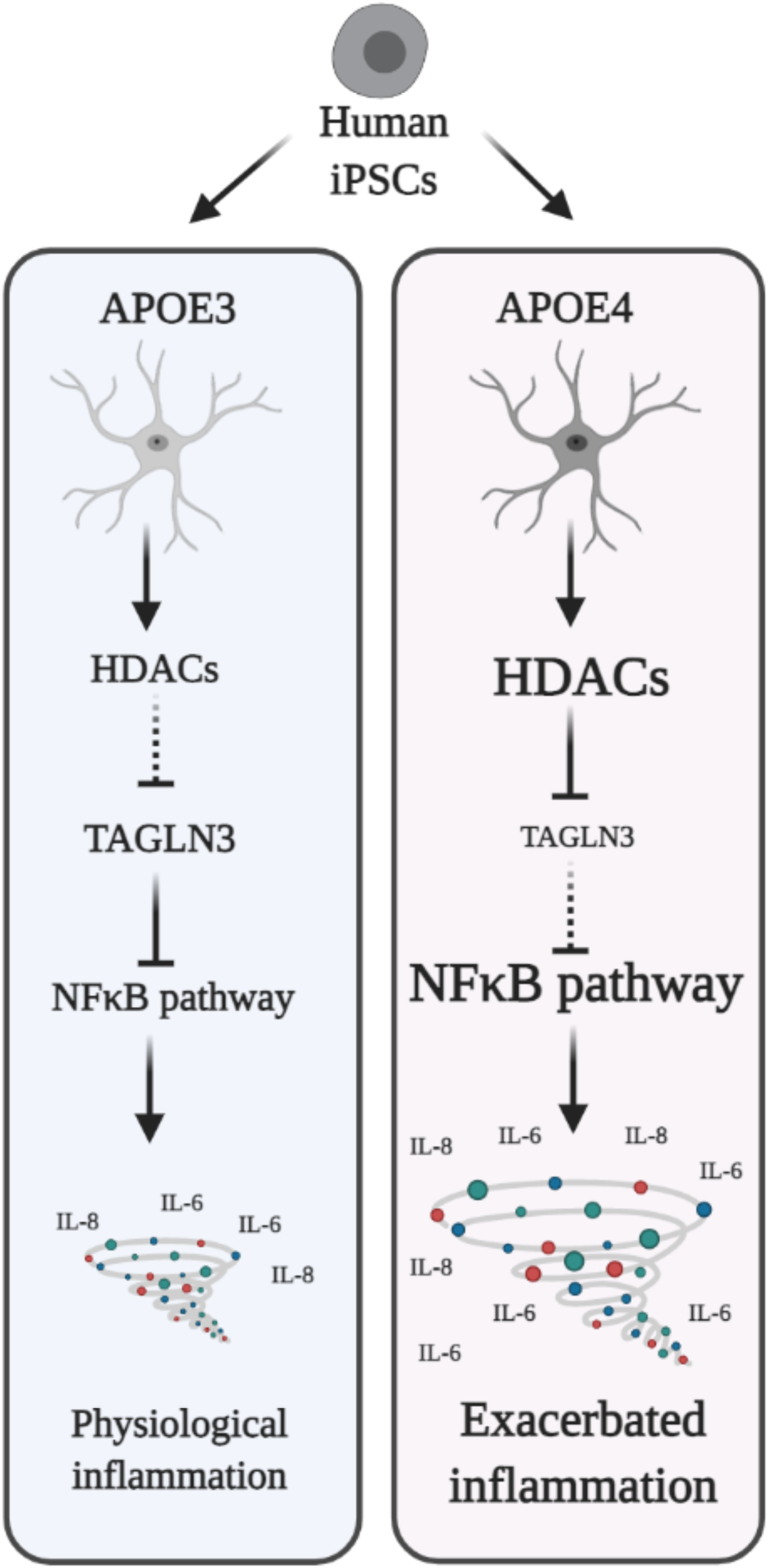

## INTRODUCTION

The Apolipoprotein E (APOE) is primarily known as a key regulator of lipid homeostasis and, within the brain, it plays a critical role in cholesterol and phospholipid transport to neurons. APOE exists in three main isoforms in human (APOE2, APOE3, APOE4) with astrocytes being the main source in the brain^1^. The *APOE4* allele is by far the major genetic variant linked to late-onset sporadic AD (sAD)^2,3^ as it is present in nearly 40% of sAD patients. APOE4 has been previously involved in Aβ plaque formation^4,5^ and Tau-mediated neurodegeneration^6^, two key hallmarks of AD. Chronic neuroinflammation is another key feature in AD and the *APOE4* allele was found to correlate with reactive gliosis in the AD brain^7^. Thus, a large body of evidence has progressively unveiled pleiotropic functions for APOE^8^, including in inflammation^9–12^. Indeed, studies have shown a cross-talk between APOE and inflammatory mediators such as cytokines^13^, revealing an intricate APOE-mediated regulation of inflammatory and immune responses. Yet, the known mechanisms by which APOE variants may affect the control of inflammation in the brain are scarce ^12^ and so, the precise role of APOE4 in astrocytes and its mechanistic connection to pathological neuroinflammation remain unclear. Identifying altered mechanisms underlying early inflammatory dysfunctions in APOE4 carriers may unveil novel APOE-mediated regulation of pathogenic mechanisms in the inflamed brain.

Neuroinflammatory changes are amongst the earliest brain manifestations in AD^14^, even before the onset of beta amyloid (Aβ) accumulation, and are increasingly considered as critical in pathogenesis initiation and progression^15–17^. Further supported by converging genetic data linking an increased AD risk to inflammatory-related genes, neuroinflammation has therefore emerged as one of the possible drivers supporting plaque formation in the brain of AD patients^18–20^. Yet, the mechanisms of pathogenic inflammation in the brain remain largely unknown. Consequently, identifying the early drivers of aberrant neuroinflammatory responses remain an urgent research need, owing the potential to identify novel therapeutic targets. To do so, however, the prevailing neuron-centric view in AD^21^, based on the dominant amyloid hypothesis^22^, has to be drifted to one that also considers the contribution of glial cells, the major mediators of neuroinflammation. Astrocytes are numerously and functionally an important glial cell type in the human brain, but were long neglected in neurodegenerative studies though gaining an increased interest^23^. Astrocytes provide homeostatic control to the brain^24^, support neuronal functions, maintain blood brain barrier integrity and also regulate innate and adaptive immune responses in the central nervous system (CNS)^25^. Morphology changes in astrocytes have long been observed in the context of neurodegenerative diseases and linked to dysregulation of inflammatory responses^26^. The critical role of reactive astrocytes in AD and their relevance as cell targets was demonstrated in AD mouse models^27,28^. Therefore, mechanistic understanding on the astrocytic pathways controlling inflammation might shed new light on the earliest events involving APOE4 in sporadic AD.

Given the differences between human and rodent astrocytes^29^, it is becoming urgent to get a more comprehensive understanding on how APOE4 could affect inflammatory responses from human astrocytes. Patient-specific iPSCs with AD-linked polymorphisms at risk for AD, together with gene-editing techniques, offer promising models for studying disease pathogenesis in relevant cell types, including human astrocytes^30,31^. Leveraging on these technologies, we found that APOE4 iPSC-derived astrocytes exhibit a higher basal expression of inflammation-related genes and an enhanced inducibility in response to cytokines when compared to their APOE3 counterparts. Furthermore, we identified a novel APOE-TAGLN3-NF-kB pathway that is dysregulated in astrocytes from AD patients carrying the APOE4 variant. Indeed, in APOE4 astrocytes, we found that TAGLN3 downregulation exacerbates inflammation through NF-kB activation. Further investigations demonstrate that such a downregulation is under the control of APOE4-dependent histone desacetylase (HDAC) activation. We showed that pharmacological intervention at different levels of this newly described regulatory axis, including TAGLN3 supplementation, can rescue the disease phenotype observed in astrocytes. Further validated in human brain samples, the description of this neuroinflammatory regulatory axis suggests TAGLN3 as a lead candidate to prevent brain damages caused by inflammatory episodes, thus providing an attractive new venue for the development of prophylactic therapeutics.

## RESULTS

### APOE4 carriers display a pro-inflammatory phenotype in patient-specific hiPSC-astrocytes

To first determine whether carrying the *APOE4* allele can affect inflammatory-associated functions in human astrocytes, we used multiple hiPSC lines derived from sAD patients and non-demented (ND) controls. Human iPSC lines presenting different *APOE* genotypes, including [*APOE3/E3*], [*APOE3/E4*] and [*APOE4/E4*] (Fig. 1a and Extended Data Fig. 1a,b; Supplementary Table 1), were differentiated into neural progenitor derivatives prior to generating astrocyte-like cells^32^ (Fig. 1a). Successful differentiation and high purity of cultures was validated by the expression of a set of astrocytic markers (*GLAST*, *GFAP*, *VIMENTIN*, *AQUA4*, *CD44*) along with the absence of cells expressing the neuronal markers *NEUN* and *TUJ1* in the culture (Fig. 1a and Extended Data Fig.1c,d). Importantly, immunoblotting analysis revealed the effective generation of APOE-producing astrocytes (Fig. 1b), a critical feature for the rest of our investigations. Differentiated astrocytes were then challenged with a cocktail of pro-inflammatory cytokines (IL-1β, MCP-1 and TNF-α), representative of the major microglia-secreted inflammatory mediators in AD^33^ (Fig. 1a). Demonstrating responsiveness and thus, astrocyte activation, gene expression profile highlighted the upregulation of four pre-selected key pro-inflammatory mediators, namely *IL6*, *IL8*, *IL1B* and *CCL2* (Fig. 1c). Most interestingly, our data highlighted significantly higher levels of these genes in an APOE4-dependent manner (Fig. 1c). Western Blotting and ELISA on IL-6 and IL-8 (not included in our cocktail), further confirmed their upregulation at the protein level. As per RNA analyses, upregulated protein levels were directly correlated to APOE4, following a rank potency order of [*APOE4/E4*] > [*APOE3/E4*] > [*APOE3/E3*] (Fig. 1d-f). Of note, for several of the analyzed inflammatory mediators, APOE4 astrocytes demonstrated higher basal cytokine levels when compared to APOE3 astrocytes (Fig. 1c). Our results suggested that the differential efficacy in the inflammatory response was dependent on APOE and not influenced by the AD status of the patient/donor as [*APOE4/E3*] astrocytes derived from a ND control showed cytokine levels similar to those obtained from sAD patients (Extended Data Fig. 1e). Supporting our observations, [*APOE3/E3*] astrocytes from a sAD patient showed an overall reduced inflammatory response compared to sAD *APOE4* carriers (Extended Data Fig. 1e). Altogether, this suggests a central role for APOE in modulating inflammation in human astrocytes.

**Fig. 1:**
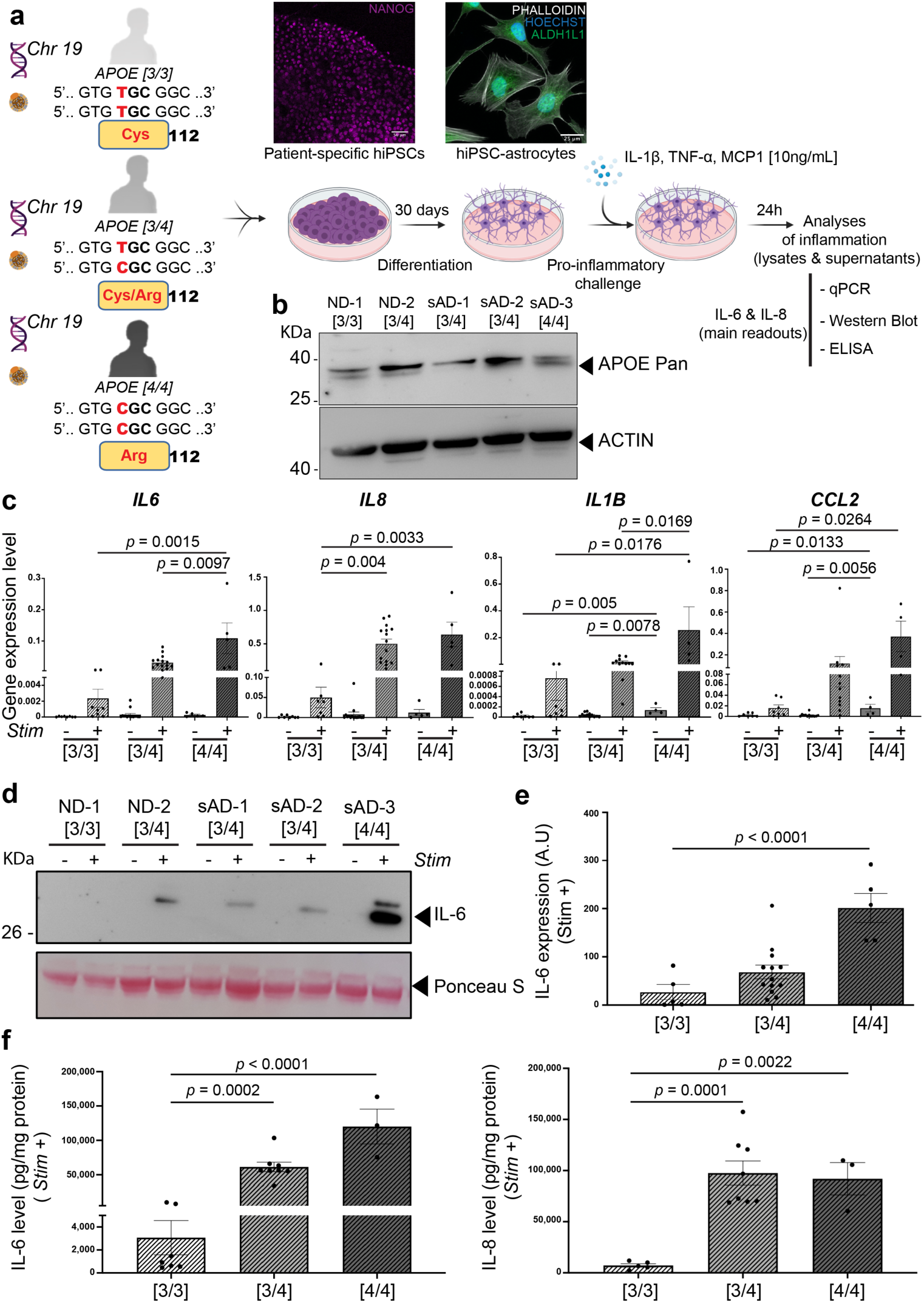
hiPSC-derived APOE4 astrocytes display an exacerbated inflammation. hiPSCs from a non-demented (ND) control donor and sporadic AD patients (sAD), carrying different *APOE* polymorphisms [*APOE3/E3*] ([3/3]), [*APOE3/E4*] ([3/4]) and [*APOE4/E4*] ([4/4]), were differentiated into astrocytes prior to challenging with a pro-inflammatory cocktail (IL-1β, MCP-1 and TNF-α). Twenty four hours after the stimulus (*Stim*), cell lysates and supernatants were analysed for inflammatory mediators. **a**) Scheme depicting the experimental paradigm. Top inserts are showing immunostainings against the NANOG pluripotency marker (purple, top left) and ALDH1L1 positive astrocytes (green, top right) counterstained with the actin filaments marker PHALLOIDIN (white) and the Hoechst blue nuclei marker (blue). **b)** Representative Western blots validating the APOE expression (lysates) in patient-specific hiPSC-astrocytes. ACTIN was used as loading control. **c)** Quantification of *IL6, IL8, IL1B* and *CCL2* mRNA levels, using qPCR analyses, from multiple patient-specific astrocytes. **d)** Representative Western blots for IL-6 production (supernatants) from patient-specific astrocyte cultures. Ponceau staining was used as loading control. **e)** Secreted levels of IL-6, quantified by relative intensities, from Western blot analyses. **f)** Absolute quantification of secreted IL-6 and IL-8 by ELISA. In all assays, one [3/3], three [3/4] and one [4/4] lines were analysed, with 2 to 7 independent batches of differentiation for each line. Data are presented as mean ± SEM using one-way ANOVA followed by *post hoc* Tukey’s test (C) or Dunnett’s test (E, F). In **e**, A.U. stands for arbitrary units (see methods). Scale bars: in **a**, 50µm (Top left insert) and 25 µm (Top right insert).

### APOE4 Knock-In and APOE knock-Out drive a pro-inflammatory phenotype in human astrocytes

When using patient-specific iPSCs as a modeling strategy to study the specific action of a given protein in human cell derivatives, one must consider the possibility that the genetic diversity from each individual could bring confounding factors on a given process. Therefore, to unequivocally investigate the role of APOE in controlling astrocytic inflammatory responses, we leverage the power of CRISPR technologies. We generated [*APOE4/E4*] Knock-In (Ki) and APOE Knock-Out (KO) isogenic clones (n=3 clones/ line) from [*APOE3/E3*] hiPSCs derived from a ND donor and a sAD patient (Fig. 2a and Extended Data Fig. 2). As before, isogenic [*APOE4/E4*] astrocytes demonstrated increased cytokine levels at both, RNA and protein levels in either unstimulated or stimulated conditions when compared to the control (Fig. 2b and Extended Data Fig. 3a-c). This reinforced the notion that APOE3 controls inflammation in astrocytes and indicated that the presence of the *APOE4* allele leads to chronic inflammation. Of further interest, we observed similar results from isogenic APOE-KO astrocytes, also displaying significantly higher basal release of pro-inflammatory mediators and greater responses upon stimulation, as measured by IL-6 and IL-8 levels (Fig. 2b and Extended Data Fig. 3a-c). Importantly, our results were highly reproducible between three independent genome-edited clones. This ruled out possible off-target effects but also the possibility that the pro-inflammatory phenotype we observed in patient-specific astrocytes was the result of a reminiscent disease state prior to reprogramming. Noteworthy, similar results were obtained from the [*APOE3/E3*] sAD line and its isogenic APOE4 and APOE-KO genome-edited clones (not shown). Together, these results indicate that the effects observed in APOE4 astrocytes could be the result, at least in part, of a loss-of-function of APOE3.

**Fig. 2:**
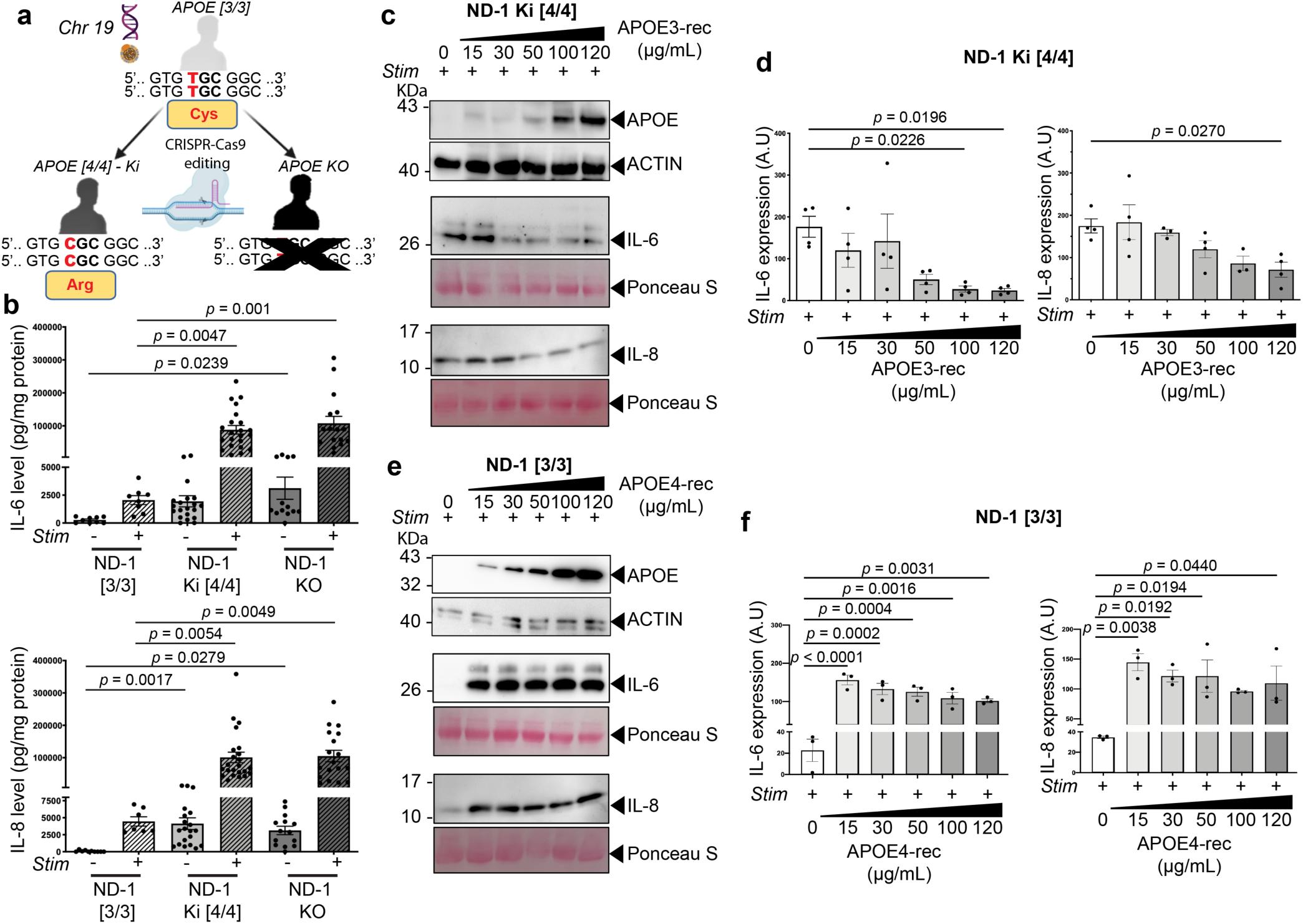
CRISPR-Cas9 editing and pharmacological interventions reveal an APOE-dependent control of inflammation in human astrocytes. **a)** Scheme depicting the experimental paradigm to generate isogenic controls, allowing the generation of [*APOE4/E4*] Knock-In (Ki) and *APOE*-knock-out (KO) hiPSCs from [*APOE3/E3*] hiPSC lines. **b)** Quantification by ELISA of the levels of IL-6 and IL-8 in the supernatants of astrocyte cultures from a non-demented donor (ND-1 [3/3]) and its respective Ki ([4/4]) and KO isogenic controls. In this assay, astrocytes from the ND-1 [APOE3/E3] line and three independent isogenic clones [*APOE4/E4*]-Ki and [*APOE*-KO] were analysed, with 4 to 9 independent batches of differentiation for each line. **c-f)** Upon application of a pro-inflammatory stimulus (Stim), APOE3 and APOE4 as recombinant proteins (APOE3- and APOE4-rec), were respectively applied on [*APOE4/E4*]-Ki or [*APOE3/E3*] astrocytes. Dose-response analyses were performed. **c,d)** Representative Western blots (**c**) and quantification (**d**) of IL-6 and IL-8 from APOE3-rec treated [*APOE4/E4*]-Ki astrocytes. **e,f)** Representative Western blots (**e**) and quantification (**f**) of IL-6 and IL-8 from APOE4-rec treated [*APOE3/E3*] astrocytes. **c-f**, astrocytes from the ND-1 [*APOE3/E3*] line and three independent isogenic clones [*APOE4/E4*]-Ki and [*APOE*-KO] were analysed, with 1 to 3 independent batches of differentiation for each line. Data are presented as mean ± SEM using one-way ANOVA followed by post hoc Dunnett’s test (**b**,**d** and **f**). In **d** and **f**, A.U. stands for arbitrary units (see methods).

To confirm that higher basal cytokine levels in astrocytes could represent a chronic pro-inflammatory phenotype and increase the response to stimuli, we assessed the daily net production of both IL-6 and IL-8 after a single stimulation with the cytokine cocktail (Extended Data Fig. 3d). IL-6 and IL-8 production was maintained at a high level over time in [*APOE4/E4*] astrocytes, whilst returning back to levels of the unstimulated condition within 5 days after stimulation in [*APOE3/E3*] astrocytes (Extended Data Fig. 3d). Overall, these data demonstrate that a single nucleotide polymorphism in *APOE* that converts *APOE3* into *APOE4* is sufficient to prime astrocytes towards a phenotype displaying chronic inflammation and exacerbated responses to a pro-inflammatory situation. Likewise, APOE-KO astrocytes phenocopied such pro-inflammatory and exacerbated response, positioning APOE3 as a core regulator of inflammation in human astrocytes and suggesting that APOE4 confers a higher susceptibility to inflammation. Thus, a pro-inflammatory environment in the brain of APOE4 patients, originating either from peripheral or central inflammatory events, might further create a positive loop in an APOE4-dependent manner in astrocytes and establish a pathogenic substrate for disease development/progression.

### APOE4-dependent pro-inflammation is attenuated by APOE4-structure correction or APOE3 supplementation

Our initial analyses in patient-specific astrocytes revealed that APOE4 homozygotes display greater inflammatory response than APOE4 heterozygotes (Fig. 1c-f). This suggested that the presence of APOE3 together with APOE4 in astrocytes could reduce, but not fully rescue, the pro-inflammatory effect of APOE4 alone. Accordingly, we next sought to determine whether exogenous APOE3 could modulate the inflammatory response from [*APOE4/E4*] astrocytes. The uptake of exogenous APOE3 by astrocytes upon supplementation in the culture media was first confirmed (Fig. 2c). Addition of recombinant APOE3 to the culture significantly reduced the release of both IL-6 and IL-8 from [*APOE4/E4*] astrocytes upon stimulation (Fig. 2c,d). Inversely, incubation of [*APOE3/E3*] astrocytes with human APOE4 recombinant protein led to an exacerbated inflammatory response upon stimulation (Fig. 2e,f).

Whereas APOE3 supplementation showed a dose-response anti-inflammatory effect in the tested range of concentrations (from 15 µg/mL to 120 µg/mL), the strong pro-inflammatory effect of APOE4 was immediate and reached a plateau from the first tested dose (*i.e.* 15 µg/mL). In line with our previous results highlighting the role of APOE independently of an AD background, these experiments further demonstrate the direct regulation of inflammation by APOE3 and the exacerbated response obtained by the presence of the APOE4 isoform.

To further validate the role of APOE4 as a direct mediator of exacerbated inflammatory responses, [*APOE4/E4*] astrocytes were incubated with the PH-002 compound, a well-characterized APOE4 structure corrector that converts APOE4 into an APOE3-like molecule^34^. We observed that PH-002 strikingly reduced the pro-inflammatory phenotype displayed by [*APOE4/E4*] astrocytes (Fig. 3a-c). These data showed that APOE3 finely tunes the inflammatory response from human astrocytes and that conformational changes in APOE4 may be sufficient to deregulate a control system that is otherwise safeguarded by APOE3. Implicit in this idea is that APOE4 may act through a dominant negative effect.

**Fig. 3:**
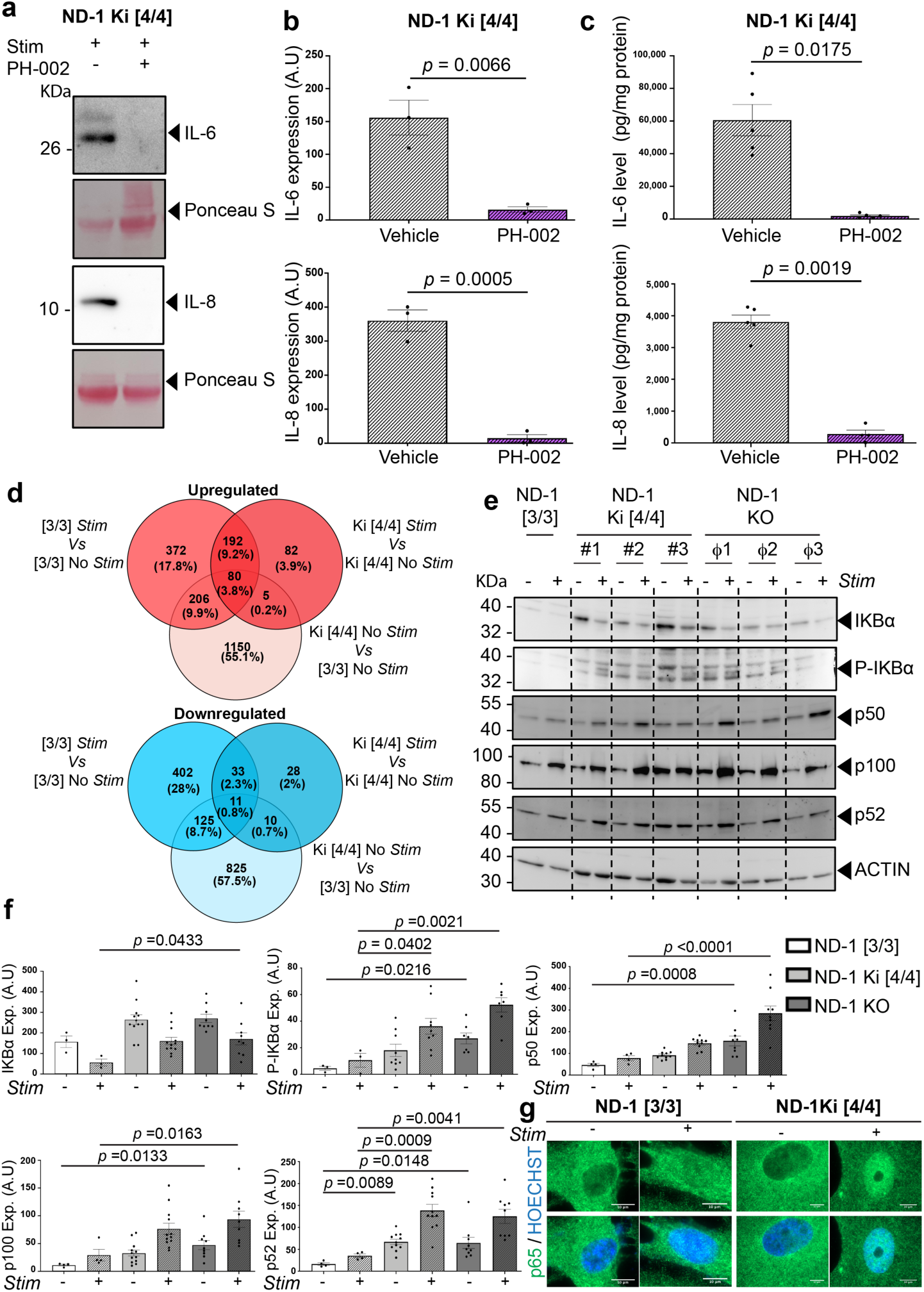
APOE4-dependent inflammation involves NF-κB hyperactivity and can be pharmacologically reversed by APOE4 structural correction. (**a-c**) Along with their stimulation with a pro-inflammatory cocktail (Stim), [*APOE4/E4*]-Ki astrocytes were treated with an APOE4 structure corrector (PH-002, 100 nM) or its control vehicle. Three independent isogenic clones [*APOE4/E4*]-Ki astrocytes were analysed with 1 to 2 independent batches of differentiation for each clone. Representative Western blots (**a**) and quantification (**b**) of IL-6 and IL-8 from PH-002 treated [*APOE4/E4*]-Ki astrocytes. (**c**) Quantification of the levels of IL-6 and IL-8 in the supernatants of [*APOE4/E4*]-Ki astrocytes treated with PH-002 24 hours prior to stimulation, as measured by ELISA. (**d**) Venn diagrams depicting the distribution of upregulated (red circles, top) and downregulated (blue circles, bottom) genes according to their differential expression between [*APOE3/E3*] and their [*APOE4/E4*]-Ki isogenic astrocytes, either in unstimulated (No stim) or stimulated (Stim) conditions. Fold changes (FC) cut-off used for analyses were ≥1.5 and ≤0.6 for 975 upregulated and downregulated genes, respectively. **e**,**f**) Representative Western blots (**e**) and quantitative analyses (**f**) for different members of the NF-κB pathway (as indicated) in [*APOE3/E3*] astrocytes (ND-1 [3/3]) and its respective isogenic [*APOE4/E4*]-Ki (ND-1 Ki[4/4]) and *APOE*-KO (ND-1 KO) counterparts. **g**) Representative images of [*APOE3/E3*] and its respective [*APOE4/E4*]-Ki astrocytes, in unstimulated (−) and stimulated (+) conditions, immunostained against p65 (green). Nuclei were counterstained with Hoechst (blue) to help the visualization of nuclear translocation. (**e-g**) astrocytes from the original ND-1 [*APOE3/E3*] line and three independent isogenic [*APOE4/E4*] and [*APOE*-KO] clones were analysed, with 3 to 6 independent batches of differentiation for each line. Data are presented as mean ± SEM using unpaired two-tailed Student’s t-Test (**b,c**) or using one-way ANOVA followed by post hoc Dunnett’s test (**f**). In (**b,f)**, A.U. stands for arbitrary units.

### APOE4 enhances basal expression of cytokine target genes towards higher NF-kB activity

Next, we sought to investigate the mechanisms by which APOE regulates inflammation in human astrocytes by identifying genes exhibiting differential expression between [*APOE4/E4*] and [*APOE3/E3*]. To this end, we used our experimental paradigm in isogenic hiPSC-astrocytes to evaluate gene expression changes that were induced by the sole T>C substitution converting *APOE3* into *APOE4*. At first, according to our criteria (see methods), we found 1441 genes exhibiting a higher basal expression in [*APOE4/E4*]-astrocytes than in [*APOE3/E3*]-astrocytes (Fig. 3d). Among them, 286 (33%) were found to be induced by our pro-inflammatory cocktail in [*APOE3/E3*]-astrocytes and 85 (23%) in stimulated [*APOE4/E4*]-astrocytes, while 272 genes (32% of the stimulated [*APOE3/E3*]- and 76% of the simulated [*APOE4/E4*]-astrocytes) were common to both genotypes. A significant number of genes eliciting a higher basal expression in [*APOE4/E4*]-astrocytes was known to be cytokine-targeted genes such as *CCL2*, *CXCL1*, *CXCL2*, *CXCL5*, *CXCL6*, *LIF*, *LY6E*, *MX1* and *WARS* (Supplementary Table 2). In addition, we observed that genes such as *REL*, *RELB*, *NFKB1*, *NFKB2* and *NFKBIA* encoding several components of the NF-kB pathway were significantly upregulated in [*APOE4/E4*] astrocytes (Supplementary Table 2). Altogether, these data suggested that APOE4 enhances inflammation *via* an increase of NF-kB activity. Moreover, we noticed that 158 genes induced in both [*APOE3/E3*] and in isogenic [*APOE4/E4*] astrocytes exhibited no difference in their basal expression. This set of genes included several known cytokine-induced genes such as C1QTNF1, CCL20, CCL5, CSF2, CXCL8, IDO1, LOXL2, MMP10, OASL, PTGS2, SP100, STAT1, IFI6, MX2 and IRF7, suggesting their regulation towards a mechanism independent of APOE.

These results prompted us to specifically analyze the effect of the APOE genotype on the NF-kB signaling pathway. Protein analyses on different NF-kB members, those found overexpressed in [*APOE4/E4*] astrocytes, confirmed gene expression data (Fig. 3e,f). More importantly, higher levels of phosphorylated IκBα, the latter being the inhibitory regulator of NF-κB when non-phosphorylated, were also observed in [*APOE4/E4*] and APOE-KO astrocytes (Fig. 3e,f). This further suggests a hyperactive NF-κB pathway in *APOE4*-carrying astrocytes, possibly due to a loss-of-function. The levels of p50 and p100, respectively members of the canonical and non-canonical pathways, were significantly increased in APOE-KO (Fig. 3e,f). Seemingly, the levels of p52, the active derivative of p100, were also significantly increased in both [*APOE4/E4*] and APOE-KO astrocytes (Fig. 3e,f). Nuclear fractionation experiments confirmed increased nuclear translocation for p50 and p52, which is known to trigger NF-κB activation, in [*APOE4/E4*] astrocytes (Extended Data Fig. 4). Immunostainings against p65 further confirmed these observations and revealed higher nuclear translocation of p65 in [*APOE4/E4*] astrocytes (Fig. 3g). Taken together, our data highlight the NF-kB signaling as a potential key driver of APOE-mediated inflammatory responses.

### APOE4-dependent downregulation of TAGLN3 triggers NF-kB activation in human astrocytes

We next used a text mining-based software^35^ to investigate the correlations between APOE, inflammation and the genes differentially expressed in [*APOE4/E4*] astrocytes, which could provide further insights on the molecular mechanisms altered by APOE4. This approach led to identify a correlation with the reduced expression of *Transgelin 3 (TAGLN3*) in [*APOE4/E4*] astrocytes, a gene encoding the TAGLN3 protein almost exclusively expressed in the CNS^36^. We focused our attention on TAGLN3 since previous reports have associated an increase in inflammation with reduced expression of the TAGLN3 close homologue SM22a/TAGLN1 in APOE null vascular smooth muscle cells (VSMCs)^37,38^. The TAGLN3 downregulation was confirmed at the proteic level. Indeed, TAGLN3 level was readily detected in APOE3 astrocytes while barely expressed in both [*APOE4/E4*] and APOE-KO isogenic astrocytes (Fig. 4a,b and Extended Data Fig. 5a). Importantly, addition of exogenous APOE3 to [*APOE4/E4*] astrocytes restored TAGLN3 levels in a dose-dependent manner (Fig. 4c,d) and was accompanied by a remarkable decrease of the IL-6 levels (Fig. 4c), the latter confirming our previous observations (Fig. 2c,d). Interestingly, similar effects were observed with the PH-002 APOE4 structural corrector, which significantly induced TAGLN3 expression in [*APOE4/E4*] astrocytes (Fig. 4e) at the same time that the levels of p50 and p52 were reduced (Fig. 4F). Altogether, these data demonstrated mechanistic connections between APOE, the regulation of TAGLN3 and NF-kB activity.

**Fig. 4:**
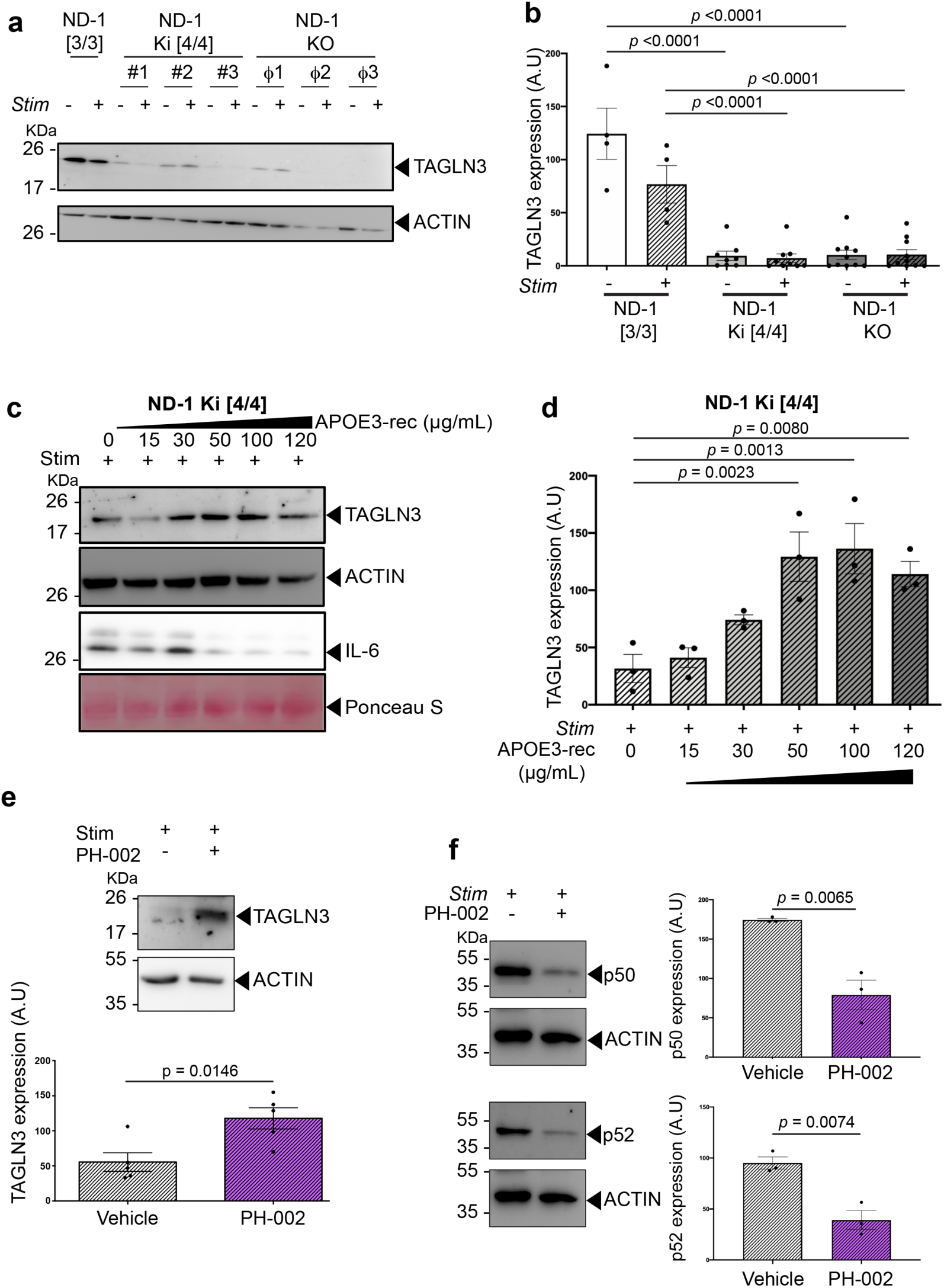
APOE4 downregulates TAGLN3 to promote pro-inflammation in human astrocytes via NF-kB activity. Representative Western blots (**a**) and quantification (**b**) of TAGLN3 from [*APOE3/E3*], [*APOE4/E4*]-Ki and [*APOE*-KO] astrocytes. **c,d**) Representative Western blots (**c**) and quantitative analyses (**d**) of TAGLN3 and IL-6 from [*APOE4/E4*]-Ki astrocytes co-treated with a pro inflammatory stimulus (Stim+) and the APOE3 recombinant protein (APOE3-rec) at different doses (n=3/dose). **e**) Representative Western blots and quantitative analyses for TAGLN3 in stimulated [*APOE4/E4*]-Ki astrocytes treated with the APOE4 structure corrector PH-002 (n=5/condition). **f**) [*APOE4/E4*]-Ki astrocytes (n=3 clones) were co-treated with a pro-inflammatory cocktail (Stim) and the APOE4 structure corrector (PH-002, 100 nM), or its control vehicle, 24 hours prior to being analysed. Representative Western blots (left panels) and quantitative analyses (right panels) for p50 and p52 levels from the lysates of the different conditions. In all different assays, astrocytes from the original ND-1 [*APOE3/E3*] line and three independent isogenic clones [*APOE4/E4*]-Ki and *APOE*-KO (when applicable) were analysed, with 1 to 3 independent batches of differentiation for each line. Data are presented as mean ±SEM using one-way ANOVA followed by post hoc Dunnett’s test (**b,d**) or using unpaired two-tailed Student’s t-Test (**e,f**). In **b**, **d**, **e** and **f**), A.U. stands for arbitrary units (see methods).

We next conducted rescue experiments to investigate whether the novel APOE-TAGLN3 axis was responsible for controlling inflammation in human astrocytes. Exogenous application of TAGLN3 reduced the inflammatory response of [*APOE4/E4*] astrocytes, as shown by significantly lower levels of IL-6 and IL-8 (Fig. 5a and Extended Data Fig. 5b-d). This was concomitant with a significant lower expression of p50 and p52 subunits in a dose-dependent manner, ultimately demonstrating for the first time that TAGLN3 controls the NF-κB pathway in human astrocytes (Fig. 5b,c). Further supporting that TAGLN3 negatively regulates inflammation in astrocytes, the stimulation of APOE3 astrocytes also led to TAGLN3 downregulation (Fig. 5d).

**Fig. 5:**
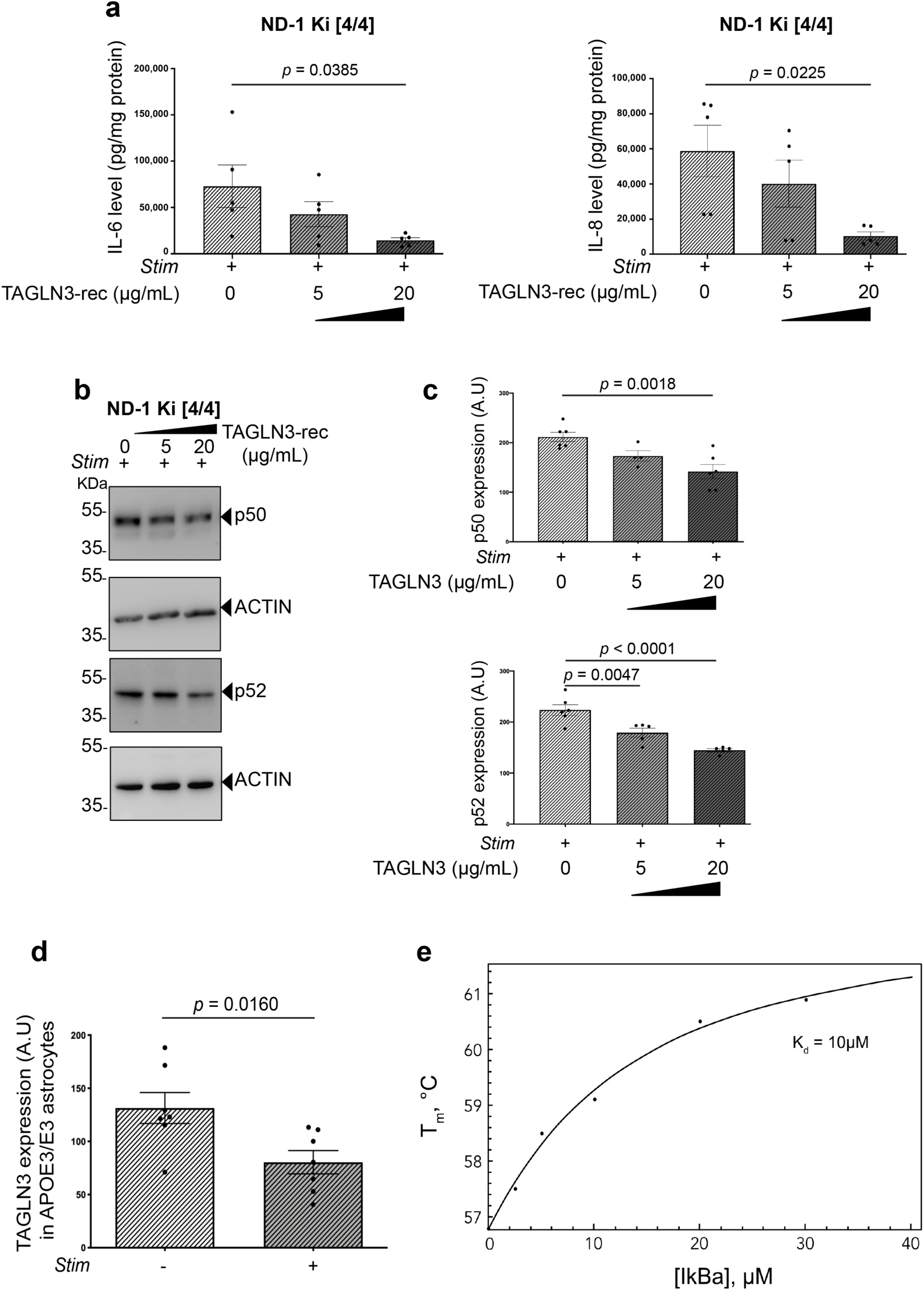
TAGLN3 exerts a control of inflammation in human astrocytes by interacting with IκBα to negatively regulate NF-κB activity. **a)** Quantification by ELISA of IL-6 and IL-8 from stimulated [*APOE4/E4*]-Ki astrocytes co-treated with two different doses of TAGLN3 (5 and 20μg/mL), as recombinant protein (TAGLN3-rec) (n=5/dose). **b,c)** Representative Western blots (**b**) and quantitative analyses (**c**) of P50 and P52 from [*APOE4/E4*]-Ki astrocytes co-treated with a pro inflammatory stimulus (Stim+) and the TAGLN3 recombinant protein (TAGLN3-rec) at different doses (n=4-6/dose). **d**) Quantification of TAGLN3 from stimulated and non-stimulated [*APOE3/E3*] astrocytes by Western blot analysis (n=7). **e**) Denaturation temperature (Tm) of TAGLN3 in the presence of increasing concentrations of IκBα (0-40 μM) was measured using nanoDSF (black dots). Data were fitted (solid line) using one-t-one binding model. In all different cell-based assays, astrocytes from the original ND-1 [*APOE3/E3*] line (**d**) and three independent isogenic clones [*APOE4/E4*]-Ki (**a,b** and **c**) were analysed, with 2 to 7 independent batches of differentiation for each line. Data are presented as mean ± SEM using one way ANOVA followed by post hoc Dunnett’s test (**a**,**c**) or using unpaired two-tailed Student’s t-Test (**d**). In **c,d** A.U. stands for arbitrary units (see methods).

Next, to investigate whether TAGLN3 regulation of NF-κB activity could be mediated, at least partly, through IκBα, we assessed TAGLN3 interaction with IκBα by following thermal denaturation of TAGLN3 in the presence of increasing concentration of IκBα using Differential Scanning Fluorimetry (nanoDSF)^39^. The progressive increase of free TAGLN3 denaturation temperature (T_m_) from 56.8°C up to 61°C for TAGLN3 in the presence of 6-fold excess of IκBα unambiguously demonstrates the existence of such interaction (Fig. 5e). Fitting obtained data using 1:1 model allowed us to estimate dissociation constant (Kd) of 10 µM. Altogether, our results identified TAGLN3 as a downstream effector of APOE and a direct regulator of the NF-kB activity through its interaction with IκBα.

### Epigenetic downregulation of TAGLN3 mediates through histone deacetylation in APOE4 astrocytes

Thereafter, we sought to identify upstream mechanisms by which the APOE4 genotype could regulate TAGLN3 expression. Previous reports have shown that SM22a/TAGLN transcription in response to TGFbeta1 is dynamically regulated by changes in the histone acetylation state^40^. Histone acetylation is regulated by histone deacetylases (HDACs) that are epigenetic regulators involved in downregulating gene expression^41^. Noticeably, previous data have shown that APOE4 increases nuclear translocation of HDACs^42^ and, moreover, several HDACs were found increased in APOE-KO mice models^43^. Our present DNA microarray array analyses revealed the upregulation of several HDACs in [*APOE4/E4*] human astrocytes such as HDAC 1/6/7/8/9/10, further confirmed by qPCR analyses for most of these HDACs (Fig. 6a and Supplementary Table 2). Thus, these observations prompted us to test whether TAGLN3 downregulation in APOE4 astrocytes could result from higher HDAC activity. [*APOE4/E4*] astrocytes were stimulated and treated with Vorinostat (SAHA), a broad inhibitor of HDAC activity. Strikingly, using such an inhibitor, we found that TAGLN3 expression was significantly increased at both genic and proteic levels (Fig. 6b,c). As expected, parallel downregulation of the IL-6 and IL-8 inflammatory mediators was observed, ruling out a global non-specific effect of HDACs inhibition (Fig. 6b-d). Thus, we concluded that APOE could regulate TAGLN3 expression, at least partly, through epigenetic regulation involving the levels of acetylation on histones.

**Fig. 6:**
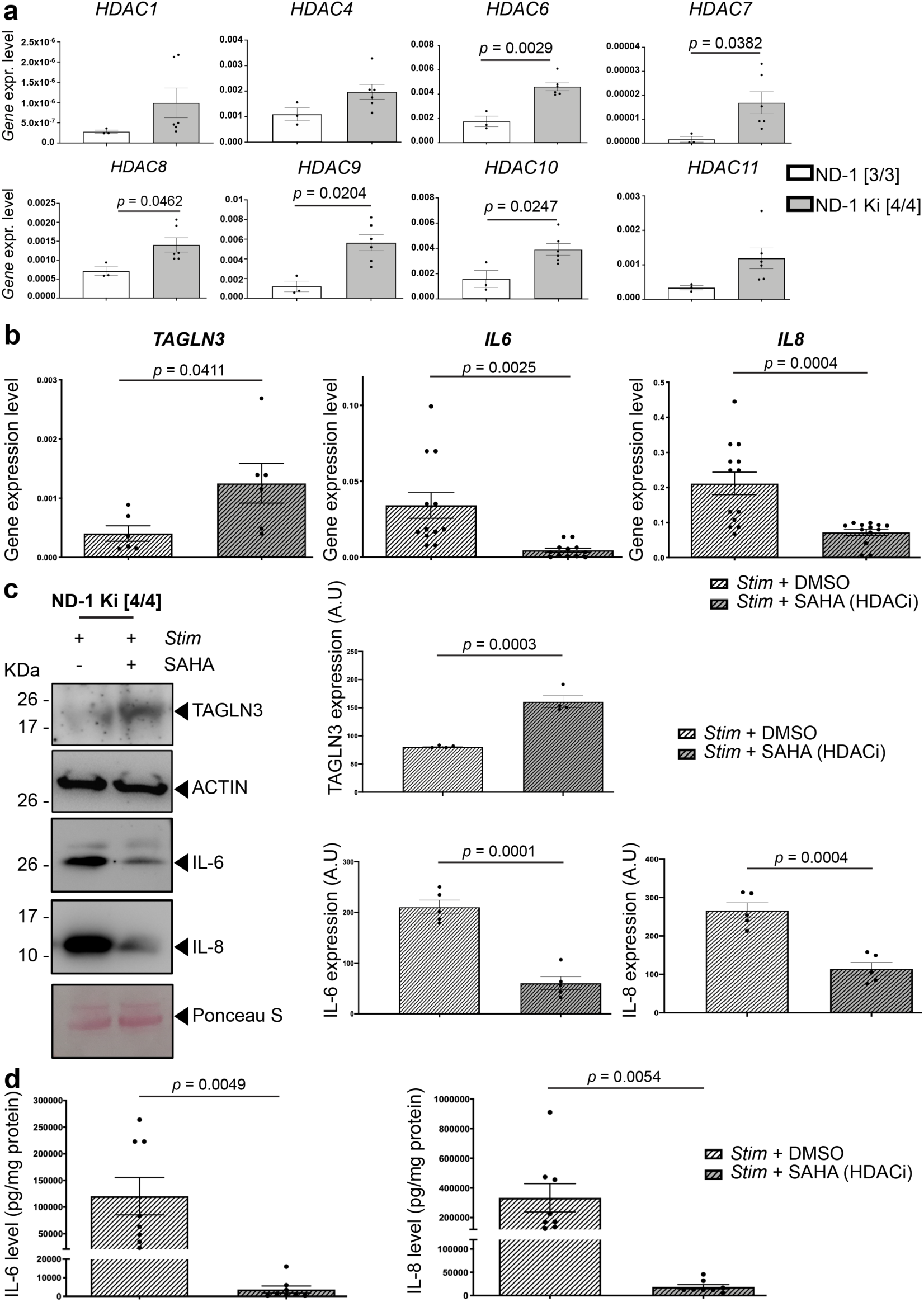
Epigenetic downregulation of *TAGLN3* results from histone hyperacetylation in APOE4 astrocytes. **a)** Gene expression levels of several HDACs (as indicated) from a non-demented [*APOE3/E3*] donor (ND-1 [3/3]) and its respective isogenic [*APOE4/E4*]-Ki (ND-1 Ki [4/4]) clones (n=3 clones). **b-d)** [*APOE4/E4*]-Ki astrocytes were treated with the broad spectrum HDAC inhibitor, SAHA or with DMSO as control. Cells were stimulated with a pro-inflammatory cocktail (Stim) and evaluated for TAGLN3, IL-6 and IL-8 expression. **b)** Bar charts showing the *TAGLN3*, *IL6* and *IL8* mRNA expression, as assessed by qPCR analysis. **c)** Representative Western blots (left panel) and quantitative analyses (right panel) for TAGLN3, IL-6 and IL-8 expression. **d**) IL-6 and IL-8 levels were assessed by ELISA from the supernatants of the different cultures/conditions. In these assays, three independent isogenic clones [*APOE4/E4*]-Ki were analysed, with 2 to 4 independent batches of differentiation for each line. Data are presented as mean ± SEM using unpaired two-tailed Student’s t-Test. In **c**, A.U. stands for arbitrary units (see methods).

### TAGLN3 downregulation is a common trait in brains from AD patients

Next, we measured TAGLN3 expression in multiple specific hiPSC lines derived from sAD patients and ND controls. We confirmed that TAGLN3 downregulation was strongly associated with the APOE4 polymorphism (Fig. 7a and Extended Data 6a-d). Last, to unequivocally evaluate TAGLN3 as a new potential target in AD, we assessed TAGLN3 expression in post-mortem human brain tissues from [*APOE3/E3*] and [*APOE4/E4*] sAD patients, as well as from the brains of [*APOE3/E3*] ND controls (Fig. 7b and Supplementary Table 3). Protein analyses revealed a significant TAGLN3 downregulation in the temporal cortex of sAD patients carrying the [*APOE4/E4*] genotype (Fig. 7c,d), ultimately confirming our initial discovery based on hiPSC-based models. Of further interest, we also found that TAGLN3 was significantly downregulated in the brain of sAD patients carrying the [*APOE3/E3*] genotype (Fig. 7C,D). The latter goes in straight line with our prior observation that, beside its chronic downregulation in APOE4 astrocytes, TAGLN3 downregulation can also occur in a pro-inflammatory context (Fig. 5d), independently of APOE4. Taken together, the pro-inflammatory environment in the brain of late-stage AD patients might lead to TAGLN3 downregulation and further contribute to exacerbated inflammation.

**Fig. 7:**
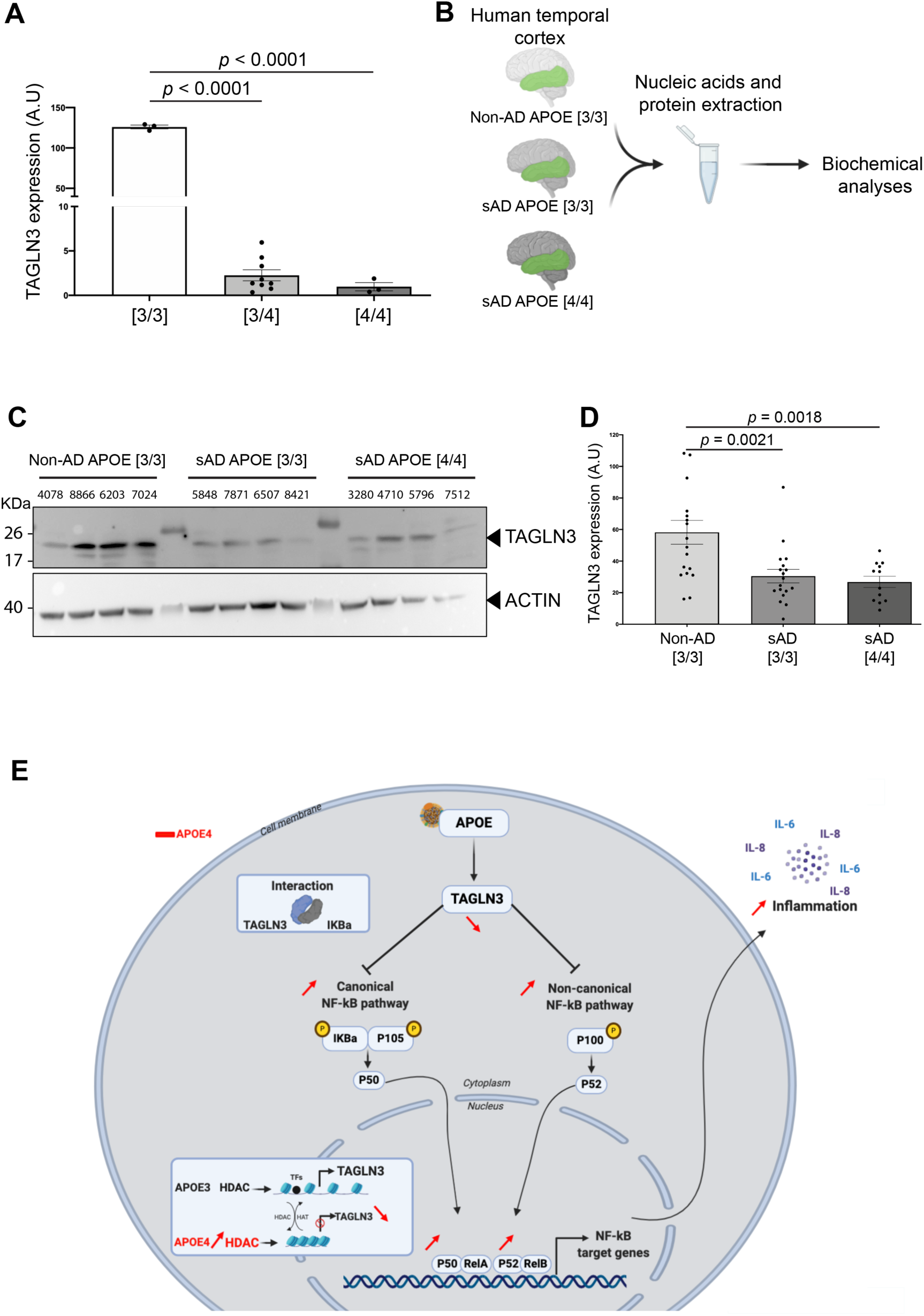
The novel APOE-TAGLN3 axis controlling inflammation in astrocytes is dysregulated in AD brains. **a**) Western blot analyses for TAGLN3 from multiple patient-specific astrocytes with different APOE genotypes. Astrocytes from one [*APOE3/E3*], three [*APOE3/E4*] and one [*APOE4/E4*] patient-specific lines were analysed, with 3 independent batches of differentiation for each line. **b**) Illustration depicting the experimental procedure used to analyse the TAGLN3 expression from post-mortem human brain samples as follows: not diagnosed for AD (Non-AD, [*APOE3/E3*] n=6) or diagnosed for AD (sAD, [*APOE3/E3*] n=6 or [*APOE4/E4*] n=4). **c,d**) Representative Western blot (**c**) and quantitative analysis (**d**) for TAGLN3 expression in the temporal cortex from the three different groups of patients as aforementioned. **e**) Schematic representation of the novel APOE-TAGLN3-NF-kB axis. Data are presented as mean ± SEM using one way ANOVA followed by post hoc Dunnett’s test. In **a** and **d**, A.U. 1000 stands for arbitrary units (see methods).

## Discussion

The APOE has been previously associated with both pro-inflammatory and anti-inflammatory responses in various pathological contexts, highlighting its intricate roles on inflammation. Several pieces of evidence have suggested that APOE4 confers a higher risk for developing CNS diseases through dysfunctions affecting the neuroinflammatory mechanisms in glial cells^12,44,45^, but the exact mechanisms remained unclear. Our present results demonstrate the key role of APOE in modulating the inflammatory response in human astrocytes and highlight mutations in *APOE*, the APOE4 variant more specifically, as a key driver of i) increased basal inflammation; and ii) exacerbated responses to cytokines stimuli that recapitulate the AD environment. Highlighting a fully novel regulatory mechanism of APOE on the NF-κB-mediated neuroinflammation (Fig. 7e), our data further support the importance of APOE in the control of inflammation. In line with recent observations that demonstrate the key role of APOE in the control of the complement cascade^12^, our results show that APOE can achieve this critical function through distinct and non-exclusive mechanisms. Based on the described repression of SM22/TAGLN expression in APOE deficient mice and the concomitant loss of its inhibitory effect on NF-kB activity^37,38^, we identified another member of the transgelin family, TAGLN3, as a novel target of APOE. Indeed, we show that the APOE4-dependent repression of TAGLN3 was responsible for the increased basal and cytokine-induced expression of inflammation-related genes. This novel mechanism could be of major importance in modulating the risk of AD since NF-κB has emerged as a regulator of aging and its over-activation was described in a number of neurodegenerative diseases^46^. Of further importance, our results revealed that the pro-inflammatory features displayed by APOE4 astrocytes were independent of other AD-associated neuronal marks such as Aβ accumulation. Thus, APOE4-dependent alterations affecting neuroinflammatory functions in astrocytes may prime the brain for neuronal-associated pathological phenotypes through chronic and/or excessive production of pro-inflammatory mediators in response to inflammatory events throughout life. Thereafter, APOE4 could also negatively impact on the worsening of the disease by other pathological mechanisms such as amyloidogenesis and tauopathy in AD, as already described^4,6^. Together, this emphasizes the necessity for a critical reconsideration of the sequence of the events underlying pathogenesis, as early glial cell dysfunctions could precede the toxic accumulation of Aβ, at least in some cases.

TAGLN3 is a member of the transgelin family preferentially expressed in the CNS, but little is known about its functions. This protein shares homology with another transgelin member, namely SM22a/TAGLN, which is abundant in VSMCs of vertebrates^47^. Although the role of TAGLN3 is much less documented than TAGLN2 and SM22a/TAGLN, they all belong to the calponin family and could be involved in the cytoskeleton organization as actin-binding proteins^48,49^. Current knowledge on TAGLN3, also known as Neuronal Protein 22 (NP22), are strictly limited to a possible role in neuronal architecture through its domains, consistent with those of cytoskeletal-interacting proteins^50,51^. A role for TAGLN3 on the astrocyte architecture cannot be excluded and will require further investigations. This is of utmost importance considering the role of the cytoskeletal machinery behind the dynamic shape changes of astrocyte morphology that drive tissue remodelling during pathogenic processes^52^. However, this is unlikely as we found TAGLN3 in the cytosol and the nucleus of astrocytes but not in the cytoskeleton (Extended Data Fig.7a), suggesting that TAGLN3 may have different functions in neurons and astrocytes. Up to now, no previous functions had ever been ascribed to TAGLN3 in astrocytes. In here, we identified a new role for TAGLN3 as a regulator of inflammation in astrocytes. The inhibitory effect of TAGLN3 on NF-kB activity can be deduced from reports highlighting the role of SM22 in its inhibition. Indeed, it was shown that SM22 deficiency promotes inflammation in VSMCs by activating canonical and noncanonical NF-kB signal pathways in vitro, respectively the degradation of IKB and p100^37,38^. The protein-ligand interaction we showed between TAGLN3 and IκBα suggest that TAGLN3 may play an important role to negatively regulate the NF-kB activity in human astrocytes.

We also identified TAGLN3 as a main protein downstream of APOE, involving epigenetic control through HDACs activity, enabling the modulation of the inflammatory responses from human astrocytes. Careful analyses of our DNA array data with our text-mining software led us to establish gene interaction maps in an attempt to postulate the mechanisms behind this epigenetic control (Extended Data Fig.7b). One of the possible ways by which APOE could exert epigenetic control of TAGLN3 can be deduced from the previously reported inhibition of SM22 in APOE deficient mice, which was shown to rely on the SP1 binding to GC element within its promoter^53^. Interestingly, SP1 was found to interact with HDACs to mediate transcriptional repression^54^. In contrast, it was demonstrated that the APOE3 genotype triggered HDAC sequestration in the cytosol^42^ and the expression of SM22 was increased by HDAC inhibitors^40^. Herein, we demonstrate that APOE4-dependent repression of TAGLN3 was triggered by the activation of HDAC activities. In line with these results, it was reported that in APOE4, nuclear translocation of HDACs led to the repressed expression of the SLC9A6/NHE6 gene^55^, repression that was also observed in our microarray data. Another non-exclusive way of epigenetic action mediated by APOE can be through the lipoprotein receptor sortilin that, by association with APOE3, facilitates the neuronal metabolism of polyunsaturated fatty acids. Interestingly, this sortilin function was disrupted by binding to APOE4^56^ and short-chain fatty acids were reported to be inhibitors of HDAC activity^57^. Thus, alteration of fatty acids metabolism resulting from sequestration of sortilin by APOE4 could enhance HDAC activity, which in turn could trigger TAGLN3 repression and the induction of inflammation. Although such hypotheses should be confirmed experimentally, our results highlight the role of APOE4 in stimulating HDAC to repress TAGLN3, alleviating its inhibitory activity on inflammation. In addition to provide an explanation for APOE4 to be an allelic risk factor in AD, our results bring new evidence on the need to consider Transgelin members as important regulators of inflammation. At present, it will be interesting to investigate whether cell type- and disease-dependent changes of expression of Transgelin members are involved in maladaptive inflammatory responses such as cytokine storms in certain sub-population of patients exposed to inflammatory events.

Our study revealed that pharmacological targeting of TAGLN3 is a valid approach to modulate neuroinflammation in astrocytes and probably a better approach as opposed to upstream modulation of APOE. Of utmost importance, combined with our *in vitro* observations, our data in human brain samples strongly support TAGLN3 downregulation as a biomarker in AD. This is especially true if we consider that TAGLN3 has already been detected in the cerebrospinal fluid^58^. Indeed, monitoring TAGLN3 downregulation in individuals could help identifying early asymptomatic stages and/or increased risk to develop the disease, as observed in *APOE4* carriers. Of further interest would be to evaluate whether TAGLN3 levels correlate with different clinical stages. In addition, developing novel TAGLN3-targeting strategies could ultimately facilitate the development of prophylactic therapies or early intervention in a stratified population at risk for numerous diseases. Such approach could lead to a first-in-class far reaching therapeutic eventually benefiting APOE4 carriers, which represent nearly 15% of the worldwide population.

## Methods

### Human iPS Cells (hiPSC)

All hiPSC lines used in this study were purchased from the Coriell biorepository. The following cell line was obtained from the NIGMS Human Genetic Cell Repository at the Coriell Institute for Medical Research [GM24666]. The following cell lines were obtained from the CIRM iPSC Repository at the Coriell Institute for Medical Research [CW70019, CW50028, CW50052, CW50069, CW50082]. Informations about hiPSC lines are indicated in Supplementary Table 1. All hiPSC lines used in this study were obtained with approved MTAs from the Aix-Marseille University thus complying with all relevant ethical regulations.

#### Culture of Human iPSC (hiPSC) lines

hiPSCs were cultured and maintained undifferentiated in a chemically defined growth media (StemMACS™ iPS-Brew XF medium; Miltenyi Biotec, Paris, France) onto growth-factor-reduced Matrigel™ (Corning)-coated plates (8,6µg/cm^2^). In short, when reaching 70–80% confluency, hiPSCs were treated with an enzyme-free solution (hereafter referred as Gentle Dissociation Solution) containing 0.5 mM EDTA (Gibco), DPBS without Ca^2+^ and Mg^2+^ (Gibco) and 1.8 mg/mL NaCl (Sigma-Aldrich). hiPSCs were incubated for 2 min in Gentle Dissociation Solution at Room Temperature (RT). The colonies were then broken into small clusters and lifted carefully using a 5 mL glass pipette at a ratio of 1:4. When necessary, differentiated areas were removed from hiPSC cultures prior to passaging in order to maintain the cultures undifferentiated before proceeding to their differentiation. hiPSC lines were maintained in an incubator (37°C, 5% CO_2_) with medium changes every day.

#### Differentiation of Neural Progenitors Cells (NPCs) from hiPSCs

Human iPSCs were differentiated into neural progenitor cells (NPCs) following a monolayer culture method with a commercial dual SMAD inhibition-mediated neural induction medium (STEMdiff™ SMADi Neural Induction Kit, Stem Cell Technologies) based on a published protocol^59^. NPCs were obtained following manufacturer’s instructions with slight modifications. Undifferentiated cultures of hiPSCs were treated with Gentle Dissociation Solution for 4 min, then the solution was removed, and cells were incubated in StemPro™ Accutase™ (Gibco) for 4 min. hiPSCs were then dislodged as single cell and transferred into Dulbecco’s Modified Eagle Medium (DMEM)/F12 (Gibco) supplemented with 20% KnockOut™ Serum Replacement (Gibco). Then, cells were centrifuged at 200 x g for 5 min, resuspended in DPBS without Ca2+ and Mg2+, counted and centrifuged once more at 200 x g for 5 min. Last, cells were resuspended in STEMdiff™ SMADi Neural Induction Kit + 10 µM Y-27632 (Tocris Bioscience), a Rho-associated protein kinase inhibitor, plated onto growth-factor-reduced Matrigel™-coated plates (8.6µg/cm2) (320,000 cells/cm2) and maintained in an incubator (37°C, 5% CO2) with medium changes every day. Between 6 and 9 days after induction, cells were harvested as single cell using Accutase™, transferred into DMEM/F12 media, centrifuged at 200 x g for 5 min, resuspended in STEMdiff™ SMADi medium + 10 µM Y-27632, plated onto growth-factor-reduced Matrigel™-coated plates (270,000 cells/cm2) and maintained in an incubator (37°C, 5% CO2) with medium changes every day for 5 additional days. Then, cells were passaged one more time following the same procedure. Five days after the last passage (i.e., between 16 to 19 days post-induction), differentiated cells were harvested as single cells using Accutase™, centrifuged at 200 x g for 5 min prior to be resuspended in STEMdiff™ Neural Progenitor Medium (Stem Cell Technologies) and seeded onto growth-factor-reduced Matrigel™-coated plates (8.6µg/cm2) at 125,000 cells/cm2. Cells were maintained in an incubator (37°C, 5% CO2) with medium changes every day and passaged with Accutase™ when reaching 80-90% confluency. Of note, hiPSC-NPCs between passage 2 and 5 were used for further maturation into astrocytes.

#### Differentiation of astrocytes-like cells from NPCs

hiPSCs-derived NPCs were differentiated into astrocytes according to a previous protocol^32^. In short, NPCs were dissociated as single cell with Accutase™ and seeded at 30,000 cells/cm^2^ on growth-factor-reduced Matrigel™-coated plates (8.6µg/cm^2^) in astrocyte medium (ScienCell, CliniSciences). Full medium changes were performed every 72 hours over a 30 day-differentiation period. When cells reached 90-95% confluency (approximately 5-7 days after initial seeding), they were passaged as single cell into astrocyte medium and cultured onto growth-factor-reduced Matrigel™-coated plates (8.6µg/cm^2^) at 30,000 cells/cm^2^. For passaging, hiPSC-astrocytes were washed with DPBS without Ca^2+^ and Mg^2+^, incubated 5 min with Accutase™, dissociated, washed with DMEM/F12 prior to be centrifugated at 200 x *g* for 5 min and resuspended with astrocyte medium. From the next passage, differentiated astrocytes were passaged at a 1:3 ratio on a weekly basis and expanded in astrocyte medium. Astrocytes between 35 and 45 days of differentiation were used throughout the study. Cells ongoing astrocyte differentiation were maintained in an incubator (37°C, 5% CO_2_) with medium changes twice a week.

#### CRISPR/Cas9 gene correction

To enable deeper analysis of APOE4 effects on inflammation in human astrocytes, we used CRISPR/Cas9 gene editing to generate [*APOE4/E4*]-Ki and APOE-KO isogenic clones (3 independent clones for each) from two different parental lines: a parental [*APOE3/E3*] line derived from a ND control donor (ND-1) and a parental [*APOE3/E3*] line derived from a sAD patient (sAD-4) (see Supplementary Table 1).

##### Preparation of the CRISPR/Cas9-ApoE sgRNA plasmid

A CRISPR/Cas9-APOE sgRNA plasmid was prepared following a published protocol^60^. Briefly, we designed a sgRNA sequence within 20 nucleotides from the target site (exon 4, amino acid 112) selected with the Bioinformatic tool CRISPOR (crispor.tefor.net). This sgRNA was chosen specifically to guide the Cas9 to cleave the double strand of DNA 3 base pairs upstream of a PAM sequence that is in close proximity to the gene region responsible for the APOE4 polymorphism. The same sgRNA was used to generate [*APOE4/E4*]-Ki and APOE-KO isogenic clones. The sgRNA sequence was as follows (5’>3”): GCGGACATGGAGGACGTGTG

We used pSpCas9 (BB)-2A-Puro (PX459) V2.0 (Addgene) and modified it in order to replace the CMV promoter by the EF1 alpha promoter. The human EF1 alpha promoter was amplified from hiPSCs with the primers pAB01 and pAB02 (5’-aattctgcagacaaatggctctagaGGCTCCGGTGCCCGTCAGTG-3’ and 5’-tccttatagtccatggtggcaccggTCACGACACCTGAAATGGAAG-3’), respectively. The final product cAB03 (*i.e.*, pX459-pEF1alpha) was then obtained by SLIC method mixing the pX459 fragment and the EF1alpha PCR product as previously described ^61^.

The sgRNA was cloned into the cAB03 open plasmid vector. For cloning, oligonucleotide pairs were hybridized with a buffer containing Tris HCl (100 mM) and NaCl (500 mM) and placed in a thermocycler ramping from 95°C to 18°C at 0.05°C/s. cAB03 was opened with BbsI (New England Labs Inc.), then ligated with annealed oligonucleotides by incubating with T4 DNA Ligase (New England Labs Inc.) 10 min at RT. The ligation product was transformed in competent bacteria DH5α (Invitrogen) following manufacturer’s instructions. After incubation for 1 h at 37°C, the bacteria were inoculated onto ampicillin-containing LB agar plates (100 µg/mL) and incubated overnight at 37°C. The next day, half of the bacterial colony was replated onto an ampicillin LB agar plate and the other half was extracted 10 min at 95°C in a lysis buffer containing 20 mM Tris HCl, 2 mM EDTA and 1% Triton 100X (pH 8). Then, a PCR was performed to verify guide insertion, using primer Pr1127 (5’-ACTATCATATGCTTACCGTAAC-3’) that binds to the backbone and the reverse oligonucleotide used to insert the sgRNA. The PCR product was put to migrate on a 2% gel to verify the insertion of the sgRNA (100 bp band) in the bacteria. One or two positive clones were then sent for sequencing to confirm the presence of the sgRNA in the plasmid after miniprep extraction (NucleoSpin® Plasmid kit, Macherey Nagel). A validated clone was then amplified, its DNA extracted by midiprep (NucleoBond® Xtra Midi EF kit, Macherey), assayed and stored at −20°C.

##### Single-Stranded OligodeoxyNucleotide (ssODN) design

We also designed single-stranded oligodeoxynucleotides (ssODN) to convert *APOE3* to *APOE4* with an editing site closed to the PAM in order to prevent recurrent Cas9 cutting in edited cells. The sequence of the donor DNA (127 nucleotides) has been chosen so that it contains the polymorphism site responsible for the *APOE4* status (bold): GCGGACATGGAGGACGTG**C**G. Arms of homology have been added on either side of this sequence to be complementary to the target region. The ssODN sequence was as follows (5’>3”): CCT GCA CCT CGC CGC GGT ACT GCA CCA GGC GGC CGC **G**CA CGT CCT CCA TGT CCG CGC CCA GCC GGG CCT GCG CCG CCT GCA GCT CCT TGG ACA GCC GTG CCC GCG TCT CCT CCG CCA CCG GGG TCA G.

##### Electroporation

Undifferentiated hiPSCs were treated with Gentle Dissociation Solution for 4 min at RT. The Gentle Dissociation Solution was then removed, and cells were treated with Accutase™ for 4 additional minutes at 37°C. Cells were collected as single cells (*i.e.*, dissociated colonies) and transferred into a tube containing a solution of DMEM/F12 supplemented with 20% of Knock-Out Serum Replacement, using a minimum of 5 times the volume of Accutase™ used for cell dissociation. Cells were centrifuged at 200 x g for 5 min, the cell pellet was washed with DPBS without Ca^2+^ and Mg^2+^. Cells were centrifuged one more time at 200 x g for 5 min. The cell pellet was then re-suspended with the Resuspension Buffer R (Neon™ Transfection System 100 µL Kit, Invitrogen) to a final concentration of 10,000 cells/µL. 100 µL of the cell suspension were transferred to a sterile 1.5 mL microcentrifuge tube and mixed with 4 µg of sgRNA (generation of KO) or 4 µg of sgRNA along with 4 µg of ssODN (1:1 ratio). Cells were then electroporated with the Neon™ Transfection System using the following parameters: 1,100V, 30 ms, 1 pulse. The electroporated cell suspension was then flushed into 1 well of a 6-well Matrigel-coated plate containing StemMACS™ iPS-Brew XF medium supplemented with 10 µM of Y-27632. Of note, one well containing cells electroporated without plasmids was used as control to define the best time window at which the puromycin treatment had to be stopped to select the electroporated cells. The next day, media was replaced with StemMACS™ iPS-Brew XF medium supplemented with puromycin (1µg/µl) until all control electroprated cells died. Puromycin-resistant cells were maintained in culture with iPS-Brew medium and changed every second day until clonal colonies could be observed.

##### Clone isolation and screening

Each individual colony (*i.e.*, those arising from a single/isolated cell after antibiotic selection) was manually picked under a microscope and cut in two halves: one half was used for genomic DNA extraction followed by PCR screening and the other half was slightly dissociated by 2-3 up and down pipetting (using a 200 µL pipette tip) and plated back onto one well of a 24-well Matrigel-coated plate containing StemMACS™ iPS-Brew XF medium supplemented with 10 µM of Y-27632. Regarding the latter, each clone was maintained in culture until (in)-validation for the KO or Ki of interest as evaluated by PCR analysis.

For genomic DNA extraction, one half of each of the puro resistant/picked colonies was individually transferred into a 0.2 ml PCR tube and centrifuged 5 min at 300 x g. The supernatant was then removed, and the pellet washed with 150 µL DPBS without Ca^2+^ and Mg^2+^ and centrifuged 5 min at 300 x g. The supernatant was removed and the cell pellet re-suspended with a 20 µL mix containing: 1X of the 5X PrimeSTAR GXL Buffer (Clontech, Takara Bio Europe), proteinase K (0.167 mg/mL, Roche Diagnostics) and ddH_2_O. DNA extraction was performed using a thermocycler with the following settings: 3 h at 55°C and 30 min at 95°C. Then, a 20 µL PCR reaction was performed using 5 µL of the extracted genomic DNA and 15 µL of a PCR mix containing 1X of the 5X PrimeSTAR GXL Buffer, PrimeSTAR GXL DNA polymerase (0.25 U/10 µL), dNTPs (200 µM each) and the specific primers for each genotypic situation reported in Supplementary Table 4; and PCR reaction was performed using the following PCR conditions: 1 min at 94°C for one cycle, then 10 sec at 98°C, 30 sec at 65°C, 45 sec at 68°C for 35 cycles and to finish, 5 min at 68°C for one cycle.

In the case of the screening of the KO clones, the PCR product was digested with the AflII enzyme (NEB) whose recognition site is at the targeted cut-off site. Thus, if a clone was KO, the enzyme was not able to cut the DNA. In the case of KI clones, PCR screening was performed by dCAPS method^62^ so that the PCR product includes a recognition site for BssHII if and only if the expected gene editing is effective. The PCR products from each clone were then digested by BssHII, and the samples were deposited to migrate onto a 3% agarose gel and revealed to analyze the size of the amplicon for each individual clone.

Expected profiles:

- APOE-KO: screening with pEN23 and pEN24 primers (digestion with AflIII)

- For APOE-KO clones: 2 bands at 576 bp and 156 bp
- For WT clones: 1 band at 700 bp
- [*APOE4/E4*]-Ki: screening with APOE03 and APOE04 primers (digestion with BssHII)

- For edition into APOE4 homozygous: 2 bands at 101 bp and 51 bp
- For WT clones (APOE3): 1 band at 152 bp

The DNA of clones identified as possibly KO or Ki was purified using magnetic beads (AMPureXP, Beckman Coulter) and the samples were sequenced by the Sanger method. Analyses of Sanger traces were done using Snapgene software. Briefly, sequences of KO or Ki PCR-based pre-selected clones were aligned with WT sequence to ultimately confirm the successful edition to generate KO or Ki isogenic clones. Positives clones were kept, amplified and confirmed again by Western blot analyses prior amplification for further experiments. A minimum of 3 independent clones for each of the edited hiPSC lines were produced to test inter-clone reproducibility of the results and so exclude biased results due to putative off-targets effects in a given clone.

### Astrocyte treatment

For each experiment/condition involving a specific treatment (as described below), a nearly confluent culture of hiPSC-derived astrocytes was passaged at a 1:3 ratio and treatment was applied when the cell reached a 70% confluence (30-45 days of differentiation).

#### Pro-inflammatory cocktail

Astrocytes were treated with a defined cocktail of pro-inflammatory mediators composed by three human recombinant proteins TNF-α, MCP-1 and IL-1β with a concentration of 10 ng/ml each for 24 hours. Each of the three proteins were purchased from Peprotech and prepared according to the manufacturer’s instructions. All three proteins were resuspended in water and so water was used as vehicle for control condition of this treatment.

#### APOE rescue experiments

For APOE rescue experiments, control [*APOE3/E3*] astrocytes or isogenic [*APOE4/E4*]-Ki clones were incubated for 24 hours with the pro-inflammatory cocktail and different doses (0, 15, 30, 50, 100 and 120 µg/ml) of APOE4 or APOE3, respectively, as human recombinant proteins (Peprotech). APOE3 was resuspended in 20 mM Sodium Phosphate, pH 7.8; 0.5 mM DTT and APOE4 was resuspended in water.

#### APOE4-structure corrector

To investigate the impact of APOE4-associated structural changes on inflammation, we incubated [*APOE4/E4*]-Ki astrocytes with 100 nM of a small molecule structure corrector for APOE4 named PH-002 ^34^ (Calbiochem, Sigma-Aldrich) during 24 hours, followed by a full medium change containing the pro-inflammatory cocktail plus a new fresh dose of PH-002 for additional 24 hours. PH-002 was resuspended in DMSO and so DMSO was used as vehicle for control condition of this treatment.

#### TAGLN3 rescue experiments

To confirm the impact of TAGLN3 on the inflammatory modulation, [*APOE4/E4*]-Ki astrocytes were incubated for 24 hours with the pro-inflammatory cocktail and different doses (0.5 and 20 µg/ml) of TAGLN3 as human recombinant protein (Abcam) prior to be analysed. TAGLN3 was resuspended 0.2% DTT, 0.32% Tris HCl, 20% Glycerol, pH 8. The same bufffer was used as vehicle for control condition of this treatment.

#### HDAC inhibition experiments

In order to study the impact of HDAC inhibition on inflammation, we incubated astrocytes with the pro-inflammatory cocktail and 10 µM of SAHA (Vorinostat, Sigma-Aldrich) as a pan-histone deacetylase (HDAC) inhibitor. SAHA was resuspended in DMSO and so DMSO was used as vehicle for control condition of this treatment.

#### Kinetic of inflammation

In order to investigate the kinetics of inflammation between [*APOE3/E3*] control and [*APOE4/E4*]-Ki astrocytes, for each genotype we set up a protocol where cells were plated at 10,000 cells/cm^2^ in 6 wells of a 6-well plate. 24 hours after seeding, cells were exposed to one dose of the pro-inflammatory mediators (10 ng/ml each, as aforementioned) during 24 hours for each well, except for one untreated control well. After 24 hours, and every 24 hours, cells from a single well were harvested (as explained below), and the medium from the other non-harvested wells was changed with fresh medium without the cocktail; thus allowing to follow the dynamic and daily net secretion of cytokines from astrocytes over a 120 hours post-stimulation period. At each time point (0, 24, 48, 72, 96 and 120 hours), supernatants were collected and stored at −80°C for further ELISA analyses, the cells were washed with DPBS without Ca^2+^ and Mg^2+^ and adherent astrocytes were lysed using RIPA buffer (Sigma-Aldrich). Lysates were then frozen at −20 °C for further protein dosage for normalization of the results from ELISA assay.

### DNA microarray

#### Sample description

For DNA microarray analyses, hiPSC-derived astrocytes obtained from the differentiation of the hiPSC ND control line (CW70019), which carries the [*APOE3/E3*] genotype, were used as control. From this hiPSC line, CRISPR-Cas9-based knock-In (KI) experiments were performed to generate three independent clones (Clones #1, #2 and #3) carrying the [*APOE4/E4*] polymorphism as aforementioned. From each of these clones, independent cultures of hiPSC-derived astrocyte-like cells were generated. Each hiPSC-derived astrocyte cultures were either treated for 24 h with the pro-inflammatory cocktail (described above) prior to harvesting the samples or placed in control untreated conditions.

#### Sample preparation

The samples were selected for microarray analyses provided that they had an >8.0 RNA integrity number, a clear gel image and no DNA contamination as shown by the histogram. Sample amplification, labelling, and hybridization essentially followed the one-colour microarray-based gene expression analysis (low input quick amp labelling) protocol (version 6.5, May 2010) recommended by Agilent Technologies. Then, 200 ng of each total RNA sample was reverse transcribed into cDNAs using the oligo dT-T7 promoter primer. Labelled cRNAs were synthesized from the cDNAs. The reaction was performed in a solution containing a dNTPs mix, cyanine 3-dCTP, and T7 RNA Polymerase and was incubated at 40°C for 2 h. RNeasy mini spin columns from Qiagen were used to purify the amplified cRNA samples. The cRNAs were quantified using a NanoDrop™ Spectrophotometer version 3.2.1 (Thermo Fisher Scientific). Six hundred nanograms of cyanine 3-labelled, linearly amplified cRNAs were used for hybridization. Hybridization on Human Gene Expression microarray slides (8×60K v2 Microarray Kit (3 slides) G4858A-039494 G3, Agilent Technologies) containing 60,000 oligonucleotide probes was performed in a 65°C hybridization oven for 17 h at 10 rpm. The hybridized microarray slides were then washed according to the manufacturer’s instructions and scanned using an Agilent Microarray Scanner with the Agilent Feature Extraction Software (Agilent Technologies). AgiND R package was used for quality control and normalization. Quantile methods and a background correction were applied for data normalization. Microarray data are available in the ArrayExpress database (164. ArrayExpress database. [www.ebi.ac.uk/arrayexpress]) under accession number E-MTAB-9625.

#### Data analysis

Only genes whose intensity values were different from background were considered. Genes were selected according to a fold change superior or equal to 1.5 for those upregulated and inferior or equal to 0.5 for those downregulated. Biological interpretation of the data was performed using the Java/Perl software Predictsearch® (Laboratoire Genex [www.laboratoire-genex.fr]), which has been previously described^35^. PredictSearch® is a powerful text mining software that identifies correlations between genes and biological processes/diseases within all scientific publications, cited at least one gene or one of its aliases in the PubMed database. The relevance of the functional correlations is supported by the Fisher test, which allows the statistical analysis of co-citations. PredictSearch® software characterizes the pathways and functional networks in which the selected genes found to be modulated are involved.

### Total RNA isolation and quantitative real-time PCR analysis for mRNA expression

The total RNA was extracted from cultured cells with NucleoSpin™ RNA Plus (Macherey-Nagel), according to the manufacturer’s recommendations. RNA concentration and purity were determined using a NanoDrop™-1000 Spectrophotometer (Thermo Fisher Scientific). Single-stranded cDNA were synthesized from 500 ng of RNA using High-Capacity RNA to cDNA™ kit (Thermo Fisher Scientific) suitable for quantitative PCR. RT-qPCR experiments were performed with the 7500 Fast Real-Time PCR System (Applied Biosystems/Life Technologies). All reactions were performed using primers ordered from Integrated DNA Technologies (IDT) as 25 nmole DNA oligos and resuspended following manufacturer’s recommendations; and the iTaq Universal SYBR Green Supermix (Bio-Rad Laboratories). Relative expression levels were determined according to the ΔΔCt method with the human housekeeping *ACTIN* gene as endogenous control for normalization. For each condition, RNA extractions were performed from independent cultures and the reported values are the mean fold change relative to the value of the control sample (untreated cell line or isogenic control line, depending of the experiment). The list of primers used in this study is reported in the Supplementary Table 5.

### Sub-cellular fractionation

Cytoplasmic and nuclear fractions were prepared from cell lysates using a ProteoExtract® Subcellular Proteome Extraction Kit (Calbiochem) according to the manufacturer’s instructions. The purity of each fraction was analyzed by immunoblotting using antibodies against Histone 3 and GAPDH for nuclear and cytoplasmic fractions, respectively.

### Immunoblotting and protein quantification

Supernatants from cells were collected and stored at −80°C in Protein LoBind Tube (Eppendorf). Cell lysates were scraped in RIPA buffer (Sigma-Aldrich) containing a cocktail of protease inhibitors (Millipore) or Protease/Phosphatase Inhibitor Cocktail (Cell Signaling) and sonicated. For each sample, the protein concentration was determined using a Bio-Rad DC™ protein assay kit (Bio-Rad). Proteins (15 µg) were loaded onto precasted Tris-Tricine 16% low molecular weight gels or pre-casted Tris-Glycine 4-20% gels (Thermo Fisher Scientific) and transferred onto nitrocellulose membranes (GE Healthcare). After blocking with 5% milk in PBS 1X, 0.2% Tween 20 (Sigma Aldrich) (or TBS for optimized protocol for detection of phospho-proteins), membranes were probed with primary antibodies against the protein of interest. Then, the appropriate HRP-conjugated secondary IgG antibodies were used (LifeTechnologies) at an appropriate dilution based on primary antibody dilution. Immunoblot signals were visualized with Essential V6 imaging platform (UVITEC) using the Amersham ECL Prime (GE Healthcare) and quantified using ImageJ software. Briefly, the expression of a given protein of interest was normalized with ACTIN for cell lysate immunoblots and with Ponceau staining for supernatant immunoblots. The list of antibodies used in this study is reported in the Supplementary Table 6.

### Differential Scanning Fluorimetry (DSF)

Thermal stability of TAGLN3 in the presence of different concentrations of NFκBα were measured in 50 mM Tris-HCl, 1 mM TCEP, buffer at pH 7.5 using nano differential scanning fluorimetry (DSF) instrument Prometheus NT.Plex (NanoTemper Technologies). NanoDSF grade capillaries were filled with 5 µM TAGLN3 solution. Concentrations of IκBα varied from 0 to 30 µM. The capillaries were loaded into the Prometheus NT.Plex instrument and the ratio of TAGLN3 fluorescence emission intensities at 330 nm (I330) and 350 nm (I350) was registered upon capillaries heating from 15°C to 95°C at rate of 1 K/min. The unfolding mid-transition temperature (T_m_) was determined from first derivative of the temperature dependence of the ratio, as implemented in Prometheus NT.Plex software. The dependence of TAGLN3 T_m_ from IκBα concentration was plotted and fitted using 1:1 binding model as described before^63^ to estimate dissociation constant (K_d_).

### Human brain tissues analysis

#### Human post-mortem brain tissues

Post-mortem human cortical brain tissue samples (temporal lobe) from 10 patients diagnosed for sAD and 6 control patients not diagnosed for AD were obtained from Neuro-CEB brain bank of Hôpital de la Pitié-Salpétrière (Paris, France) after informed consent of all participants during their life and/or their legal guardians. Characteristics of the population of patients such as *APOE* genotype, sex, age, post-mortem delay of storage and Braak stage are scoring by members of the Neuro-Ceb and listed in the **Supplementary Table 6**. All tissues were obtained following informed consent, flash frozen at time of autopsy indicated in the Supplementary Table 3 and stored at −80°C until being processed for analyses. All brain samples used in this study were obtained with approved MTAs from the Aix-Marseille University thus complying with all relevant ethical regulations.

#### Sample preparation

For protein extraction, brain tissue samples of 100 ± 5 mg were lysed in 1 mL of RIPA containing a cocktail of protease inhibitors (Millipore), homogenized with Dounce homogenizer (Dutcher), tubes were agitated 10 min at 4°C and centrifugated at 12,000 x *g* for 10 min at 4°C. Then, the supernatants were transferred into a new set of tubes and protein concentration was determined using Bio-Rad DC™ protein assay kit (Bio-Rad). Western blot protocol is the same as described before. Note that 30 µg of total protein was loaded in this case.

For RNA extraction, 80 ± 5 mg of brain tissue samples were lysed in 1 mL of TRIzol™ Reagent (Thermo Fisher), according to the manufacturer’s recommendations. RT-qPCR protocol is the same as described before.

### Immunofluorescence labelling experiments

Cells were fixed in 4% paraformaldehyde (Antigenfix, Diapath) for 10 minutes at RT before immunostaining. After blocking with PBS 1X, 3% BSA, 0.1% Triton, cells were incubated overnight at 4°C with the following primary antibodies: NANOG, ALDH1L1, Vimentin and GLAST. The corresponding secondary antibodies (coupled to Alexa Fluor® 488 or 568, LifeTechnologies, 1:800) and Hoechst (Sigma-Aldrich, 1/800) for 2 hours at RT. The list of antibodies used in this study is reported in the Supplementary Table 6. Cells were examined using a Zeiss LSM 710 Laser Scanning Confocal Microscope (Zeiss) or an Axio-Observer upright epifluorescence microscope (Zeiss) and post-acquisition was performed using the Zen software (Zeiss) and/or the Fiji (ImageJ) software.

### Quantification and statistical analysis

Sample size was chosen to ensure sufficient statistical power for each assay throughout the study. All statistical analyses were performed using GraphPad Prism software on mean values calculated from the averages of biological and/or technical replicates. In all assays, we used multiple and independent batches of differentiation from each of the tested lines/clones, adding further statistical power. In short, hiPSC lines derived from ND and sAD patients (ND-1, ND-2, sAD-1, sAD-2, sAD-3 and sAD-4) were differentiated in astrocyte-like cells with 3 to 7 independent batches for analyses as replicates. For isogenic Ki and KO clones from both ND-1 and sAD-4, 3 independent clones were generated and differentiated into astrocyte-like cells with 3 to 8 independent batches for analyses as replicates. For each line, 3 independent clones were generated to: i) avoid biased results due to off-target effect, as the chance to get the same off-target modification in 3 independent clones are statistically extremely low; ii) offer the possibility to repeat experiments in three different clones, providing enough statistical power.

Statistical significance was calculated by two-tailed unpaired Student’s *t*-test to compare two experimental groups or one-way ANOVA (multicomparison, three or more conditions) followed by Dunnett’s post-hoc test for differences of means between each group and the control group of data or followed by Tukey’s post-hoc test for multiple comparison between groups, with *P* < 0.05 considered statistically significant. The exact *P* values are always reported, except when *P* < 0.0001. No values were excluded from the analyses. Sample sizes for all experiments were chosen based on previous experiences. Independent experiments were pooled and analysed together whenever possible as detailed in figure legends. All graphs show mean values ± SEM.

## Data availability

Microarray data that support the findings of this study have been deposited in the ArrayExpress database (164. ArrayExpress database. [www.ebi.ac.uk/arrayexpress]) under accession number E-MTAB-9625.

## Acknowledgements

This work was supported by fundings from the CNRS and Aix Marseille Université. The work was also supported by the DHUNE center of excellence and a CoEN grant to S.R., and E.N., by the “FONDATION ALZHEIMER” to S.R., and E.N., by “FONDATION VAINCRE ALZHEIMER” FR-19052 to E.N., by a grant from “La Fondation NRJ-Institut de France” to E.N. L.A., was supported by the CoEN grant. LGG was supported by an ANR grant (MAD5) to S.R., and also granted by the Fondation pour la Recherche Médicale FRM FDT201904008423 and by a Fondation Vaincre Alzheimer international travel grant. This work has been carried out thanks to the support of the A*Midex foundation and the French National Research Agency funded by the French Government « Investissements d’Avenir » program (NeuroSchool, nEURo*AMU, ANR-17-EURE-0029 grant). All human brain samples used in the study were kindly provided by the Brainbank Neuro-CEB Neuropathology Network*. We thank Dr Aurélie Tchoghandjian Auphan and Thomas Perez for providing us with antibodies against members of the NF-κB pathway and the SAHA compound, respectively. We thank the Platform NeuroTimone (PFNT) of the Aix-Marseille University, especially the Stem Cell Center NeuroTimone (SCeNT) and the Interactome Timone Platform (PINT), for technical supports regarding to hiPSC experiments and nanoDSF experiments, respectively. Schematic illustration used in this study are unique and were created from the BioRender.com website.

*The NeuroCEB Neuropathology network includes: Dr Franck Letournel (CHU Angers), Dr Marie-Laure Martin-Négrier (CHU Bordeaux), Pr Françoise Chapon (CHU Caen), Pr Catherine Godfraind (CHU Clermont-Ferrand), Pr Claude-Alain Maurage (CHU Lille), Dr Vincent Deramecourt (CHU Lille), Dr David Meyronnet (CHU Lyon), Dr Nathalie Streichenberger (CHU Lyon), Dr André Maues de Paula (CHU Marseille), Pr Valérie Rigau (CHU Montpellier), Dr Fanny Vandenbos-Burel (Nice), Pr Charles Duyckaerts (CHU PS Paris), Pr Danielle Seilhean (CHU PS, Paris), Dr Susana Boluda (CHU PS, Paris), Dr Isabelle Plu (CHU PS, Paris), Dr Serge Milin (CHU Poitiers), Dr Dan Christian Chiforeanu (CHU Rennes), Pr Annie Laquerrière (CHU Rouen), Dr Béatrice Lannes (CHU Strasbourg).

## Author Contributions

L.A., and E.N. designed and conceived the whole study. P.B., J.C.I.B., I.S.M., K.B., and S.R. advised on some experiments. L.A., L.G., and D.S. performed all experiments, collected and analyzed the data. L.A., K.B., S.R., and E.N. interpreted the data. P.B. performed DNA microarray analyses and interpreted the microarray data. A.J., performed cell culture experiments, Western blot analyses and image acquisitions. L.A., L.G., L.G.G., and N.J. contributed to CRISPR-Cas9 design and plasmid constructions. F.D., and P.O.T. performed nanoDSF experiments. J.C.I.B., advised and provided training for CRISPR-Cas9 experiments. E.N., wrote the original manuscript. L.A., P.B., L.G., I.S.M., J.C.I.B., S.R., and E.N. assisted with review and editing of the manuscript. E.N. supervised all studies and manuscript preparation. All authors reviewed the manuscript and approved the final version.

## Materials & Correspondence

The materials generated and/or analysed during the current study are available from the corresponding author on reasonable request and based on feasibility.

## SUPPLEMENTARY TABLES (1 to 6)

**Table 1.** hiPSC lines used in the study.

| Line | Diagnosis | Nomenclature | Gender | Age (at sampling) | Age at onset of dementia | APOE status |
| --- | --- | --- | --- | --- | --- | --- |
| CW70019 | Non-demented control | ND-1 | Female | 67 | N/A | $\epsilon 3/\epsilon 3$ |
| CW50028 | Non-demented control | ND-2 | Male | 68 | N/A | $\epsilon 3/\epsilon 4$ |
| CW50069 | Sporadic AD | sAD-1 | Female | 66 | 64 | $\epsilon 3/\epsilon 4$ |
| CW50082 | Sporadic AD | sAD-2 | Male | 65 | 65 | $\epsilon 3/\epsilon 4$ |
| CW50052 | Sporadic AD | sAD-3 | Male | 87 | 86 | $\epsilon 4/\epsilon 4$ |
| GM24666 | Sporadic AD | sAD-4 | Male | 83 | | $\epsilon 3/\epsilon 3$ |

**Table 2.** List of selected genes found dysregulated by DNA microarray analyses. Fold change values are written in red (over-expressed genes) or in blue (under-expressed genes).

| GeneName | ND-1 Ki [4/4] #1 *No Stim*<br>*Vs ND-1 [3/3] No stim* | ND-1 Ki [4/4] #2 *No Stim*<br>*Vs ND-1 [3/3] No stim* | ND-1 Ki [4/4] #3 *No Stim*<br>*Vs ND-1 [3/3] No stim* | ND-1 [3/3] *Stim*<br>*Vs ND-1 [3/3] No Stim* | ND-1 Ki [4/4] #1 *Stim*<br>*Vs ND-1 Ki [4/4] #1 No stim* | ND-1 Ki [4/4] #2 *Stim*<br>*Vs ND-1 Ki [4/4] #2 No stim* | ND-1 Ki [4/4] #3 *Stim*<br>*Vs ND-1 Ki [4/4] #3 No stim* |
| --- | --- | --- | --- | --- | --- | --- | --- |
| CFLAR | 3,1 | 2,0 | 1,5 | 3,0 | 1,6 | 1,9 | 2,5 |
| LIF | 1,7 | 2,4 | 6,1 | 18,3 | 5,6 | 8,7 | 5,1 |
| CCL2 | 8,7 | 19,2 | 28,1 | 103,6 | 11,3 | 5,2 | 3,9 |
| CXCL1 | 6,6 | 52,2 | 15,0 | 752,0 | 82,4 | 14,3 | 39,0 |
| CXCL2 | 7,2 | 113,8 | 43,7 | 944,2 | 43,7 | 6,5 | 28,7 |
| CXCL5 | 1,6 | 24,8 | 2,8 | 268,0 | 35,5 | 7,3 | 20,4 |
| CXCL6 | 6,4 | 11,1 | 4,1 | 541,8 | 29,4 | 22,3 | 54,9 |
| NFKB1 | 2,2 | 3,8 | 1,7 | 10,2 | 3,4 | 3,3 | 5,8 |
| NFKB2 | 4,2 | 4,3 | 3,1 | 9,4 | 6,0 | 8,1 | 13,4 |
| NFKBIA | 2,0 | 2,9 | 2,8 | 13,9 | 4,8 | 4,4 | 5,3 |
| REL | 1,7 | 1,7 | 2,0 | 2,8 | 2,3 | 2,5 | 3,9 |
| RELB | 4,4 | 7,2 | 7,1 | 18,6 | 3,7 | 4,8 | 4,2 |
| TNFAIP2 | 19,5 | 87,5 | 15,4 | 275,4 | 11,8 | 4,3 | 24,5 |
| TNFAIP3 | 4,4 | 5,4 | 5,5 | 61,4 | 10,0 | 10,5 | 19,2 |
| MMP9 | 2,7 | 10,2 | 1,7 | 35,1 | 67,0 | 26,8 | 24,7 |
| ADM | 3,6 | 11,4 | 2,8 | 1,3 | 1,0 | 0,6 | 0,8 |
| RAMP1 | 10,1 | 6,1 | 4,0 | 0,6 | 0,4 | 0,4 | 0,4 |
| HDAC1 | 4,6 | 1,9 | 4,1 | 1,1 | 0,9 | 1,0 | 0,9 |
| HDAC10 | 3,4 | 2,7 | 1,9 | 0,6 | 1,1 | 1,0 | 1,5 |
| HDAC6 | 1,9 | 1,7 | 2,2 | 0,8 | 1,1 | 0,9 | 1,1 |
| HDAC7 | 1,8 | 1,5 | 1,5 | 0,8 | 1,0 | 1,1 | 1,7 |
| HDAC9 | 5,8 | 1,8 | 5,2 | 1,9 | 0,9 | 2,0 | 1,6 |
| TAGLN3 | 0,0 | 0,2 | 0,2 | 0,8 | 1,0 | 0,8 | 0,6 |
| SLC9A6 | 0,4 | 0,4 | 0,6 | 0,9 | 0,8 | 0,9 | 1,1 |
| APOE | 0,2 | 0,1 | 0,0 | 1,7 | 0,8 | 1,7 | 0,9 |

**Table 3.** Information regarding human brain tissue samples used in the study.

| Patients | Gender | Age (years old) | Braak stage | Post-mortem delay<br>(hours) | APOE status |
| --- | --- | --- | --- | --- | --- |
| Control | F | 92 | Stage I like | 21 | ε3/ε3 |
| Control | M | 73 | N/A | 10 | ε3/ε3 |
| Control | M | 84 | N/A | 15.30 | ε3/ε3 |
| Control | M | 78 | N/A | 23 | ε3/ε3 |
| Control | F | 76 | Stage IV like | 28 | ε3/ε3 |
| Control | M | 70 | Stage III like | 31 | ε3/ε3 |
| AD | F | 91 | VI | 26 | ε3/ε3 |
| AD | M | 71 | VI | 6 | ε3/ε3 |
| AD | M | 91 | VI | 11 | ε3/ε3 |
| AD | F | 85 | VI | nd | ε3/ε3 |
| AD | F | 83 | VI | nd | ε3/ε3 |
| AD | M | 84 | VI | nd | ε3/ε3 |
| AD | F | 91 | VI | 13 | ε4/ε4 |
| AD | M | 77 | VI | nd | ε4/ε4 |
| AD | M | 70 | VI | 6 | ε4/ε4 |
| AD | F | 82 | VI | nd | ε4/ε4 |

**Table 4.**
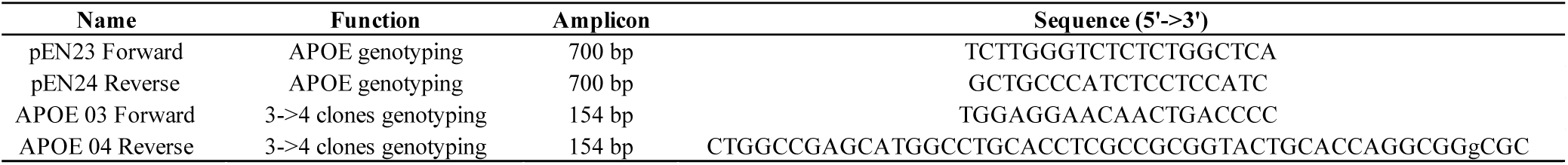
List of primers for PCR screening.

**Table 5.**
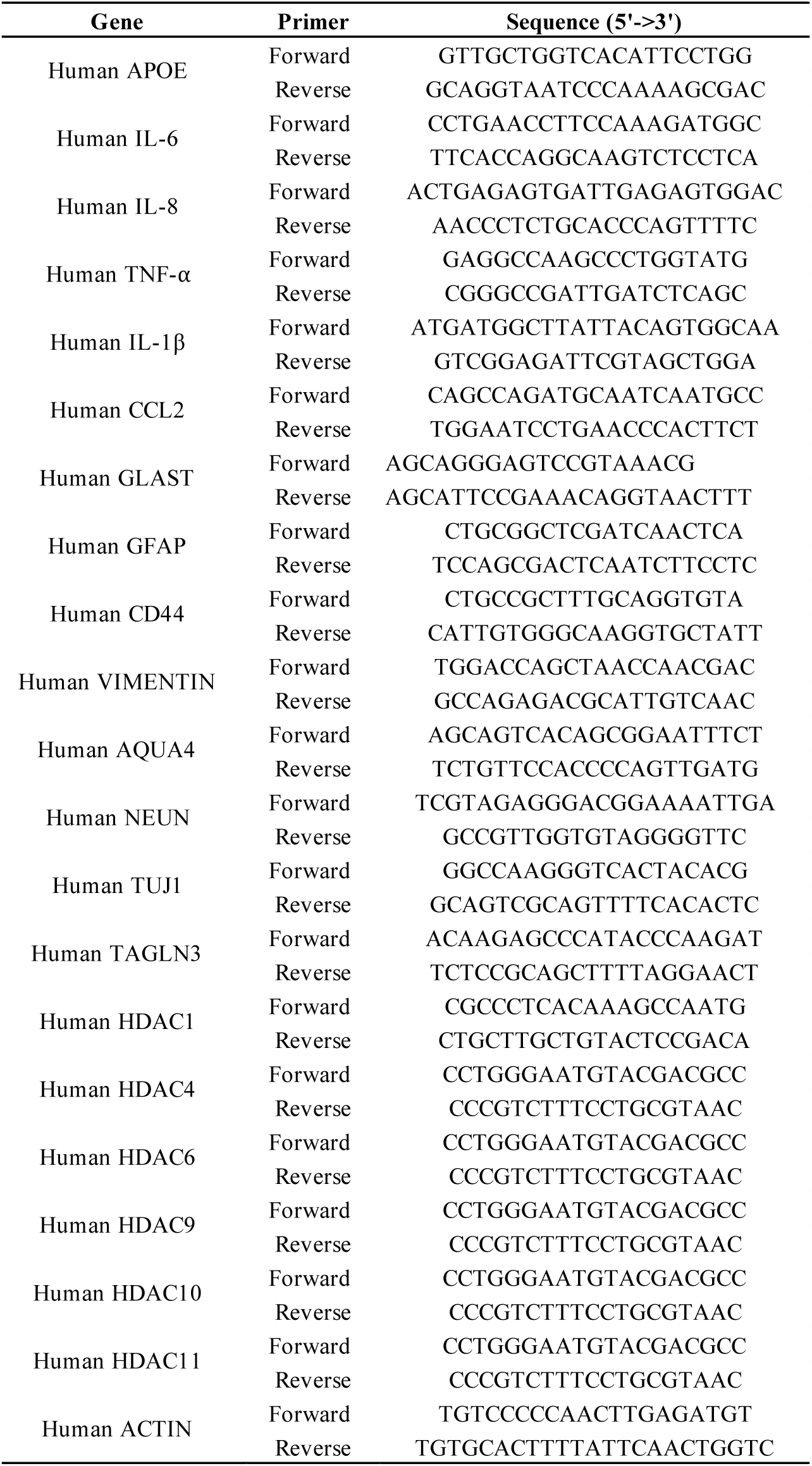
List of primers used for qPCR experiments.

**Table 6.** List of primary antibodies used for Western Blot analysis and Immunocytochemistry.

| **Antibody** | **Supplier** | **Reference** | **Dilution used** |
| --- | --- | --- | --- |
| APOE4 (4E4) | Enzo Life | ENZ-ABS295-0100 | 1/500 |
| APOE pan (F-9) | Santa Cruz | sc-390925 | 1/200 |
| IL-6 | Santa Cruz | sc-130326 | 1/200 |
| IL-8 | Santa Cruz | sc-376750 | 1/200 |
| IκBα (44D4) | Cell Signaling | 4812S | 1/500 |
| Phospho-IκBα (Ser32) (14D4) | Cell Signaling | 2859S | 1/500 |
| NFκB p50 (H-119) | Santa Cruz | sc-7178 | 1/500 |
| NFκp52 (05-361) | Millipore | 05-361 | 1/500 |
| NFκp65 (F-6) | Santa Cruz | sc-8008 | 1/200 |
| Phospho-NFκp65 (S536) | Cell Signaling | 3031S | 1/1000 |
| Actin | Sigma-Aldrich | A5316 | 1/5000 |
| GAPDH | Sigma-Aldrich | G8795 | 1/10000 |
| Histone 3 | Cell Signaling | 9715 | 1/1000 |
| NANOG | abcam | 21624 | 1/400 |
| Vimentin | Cell Signaling | 3932S | 1/200 |
| GLAST | ThermoFisher | PA5-34198 | 1/50 |
| ALDH1L1 | Abcam | ab190298 | 1/200 |

## EXTENDED DATA FILE (Arnaud et al.) (Contains 7 figure legends, 7 Extended DATA Figures)

**Extended Data Figure 1.**
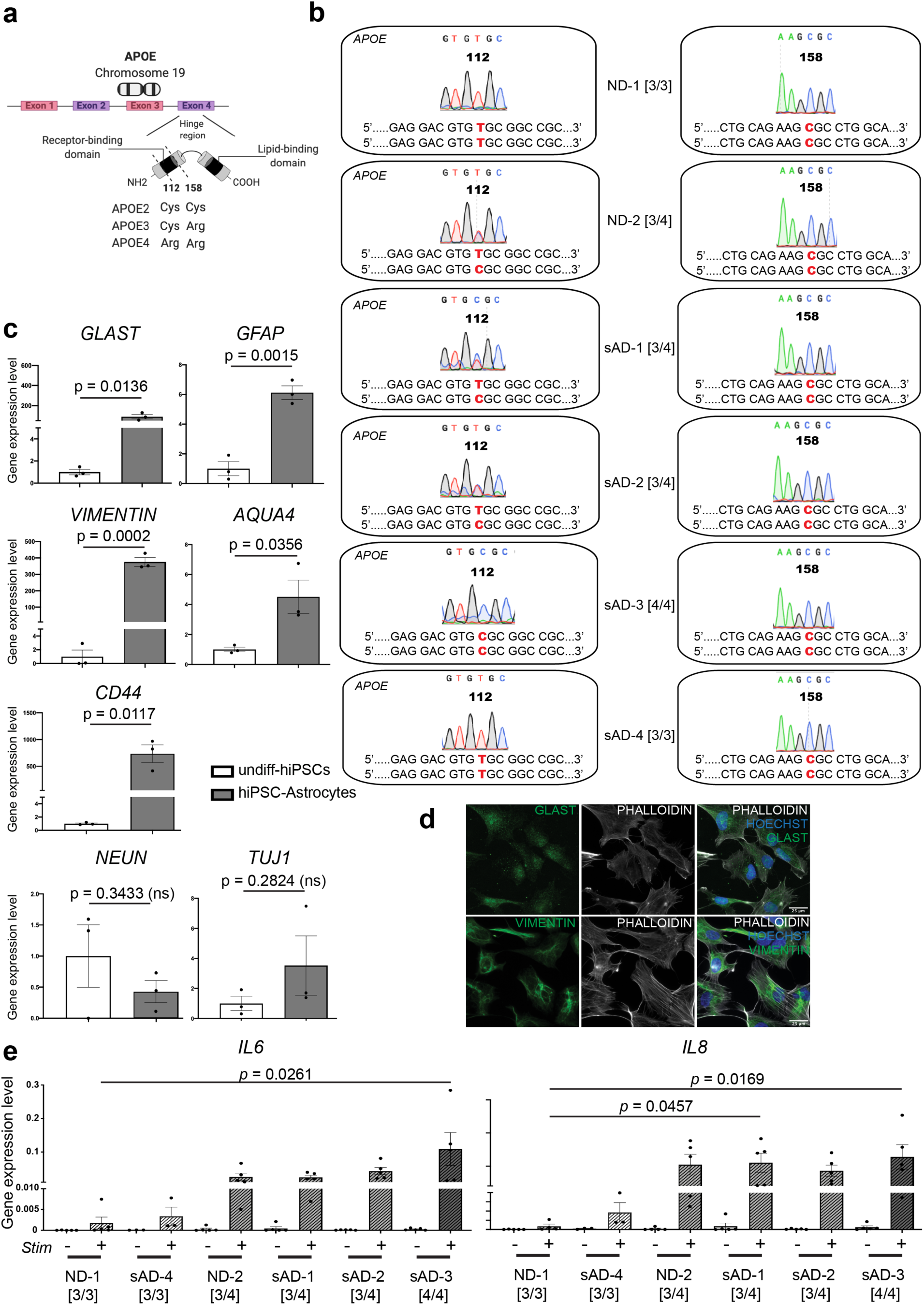
APOE polymorphism from patient-specific hiPSCs and differentiation into astrocytes with a pro-inflammatory signature driven by APOE4. **a**) Illustration representing the human APOE gene, located in the Chromosome 19. The three main isoforms for APOE – i.e., APOE2, APOE3 and APOE4 – are only different from a single nucleotide polymorphism (APOE3>APOE2 or APOE3>APOE4) or two nucleotides substitutions (APOE2>APOE4) on codon 112 and/or 158. The APOE4 isoform results from a T>C substitution that leads to an Arginine in the 112th position instead of a cysteine for the APOE3 isoform. **b**), the *APOE* genotype of hiPSCs from non-demented (ND) controls and sporadic AD (sAD) patients was given from the original source and confirmed by Sanger sequencing. Analyses were focused on the positions 112 and 158. **c**) Multiple hiPSC lines were differentiated into astrocytes (hiPSC-astrocytes), within 30 days, prior to being analyzed for astrocytic markers. The bar chart shows gene expression changes, measured by qPCR analyses, for astrocytic markers (GLAST, GFAP, CD44, VIMENTIN, AQUA4) and neuronal markers (NEUN, TUJ1), data are normalized to their respective undifferentiated hiPSCs (undiff-hiPSCs). Data are presented as mean ± SEM (n=3) using unpaired two-tailed Student’s t-Test. **d**) Representative images of hiPSC-derived astrocytes immunostained with antibodies against the human astrocyte markers GLAST (green, top panels) or VIMENTIN (green, bottom panel). Cells were counterstained with the actin filaments marker PHALLOIDIN (white) and the Hoechst blue nuclei marker (blue). **e**) hiPSCs from nondemented (ND) control donors and sporadic AD (sAD) patients, presenting different APOE polymorphisms [*APOE3/E3*], [*APOE3/E4*] and [*APOE4/E4*], were differentiated into astrocyte-like cells prior to challenging with a defined pro-inflammatory cocktail (IL-1β, MCP-1 and TNF-α, referred as Stim). The bar charts show the gene expression level for IL6 (left) and IL8 (right) in patient-specific astrocytes for each of the indicated line. In this assay, two [3/3], three [3/4] and one [4/4] lines were analysed, with 3 to 5 independent batches of differentiation for each line. ns stands for non-significant. Scale bars in d, 25µm.

**Extended Data Figure 2.**
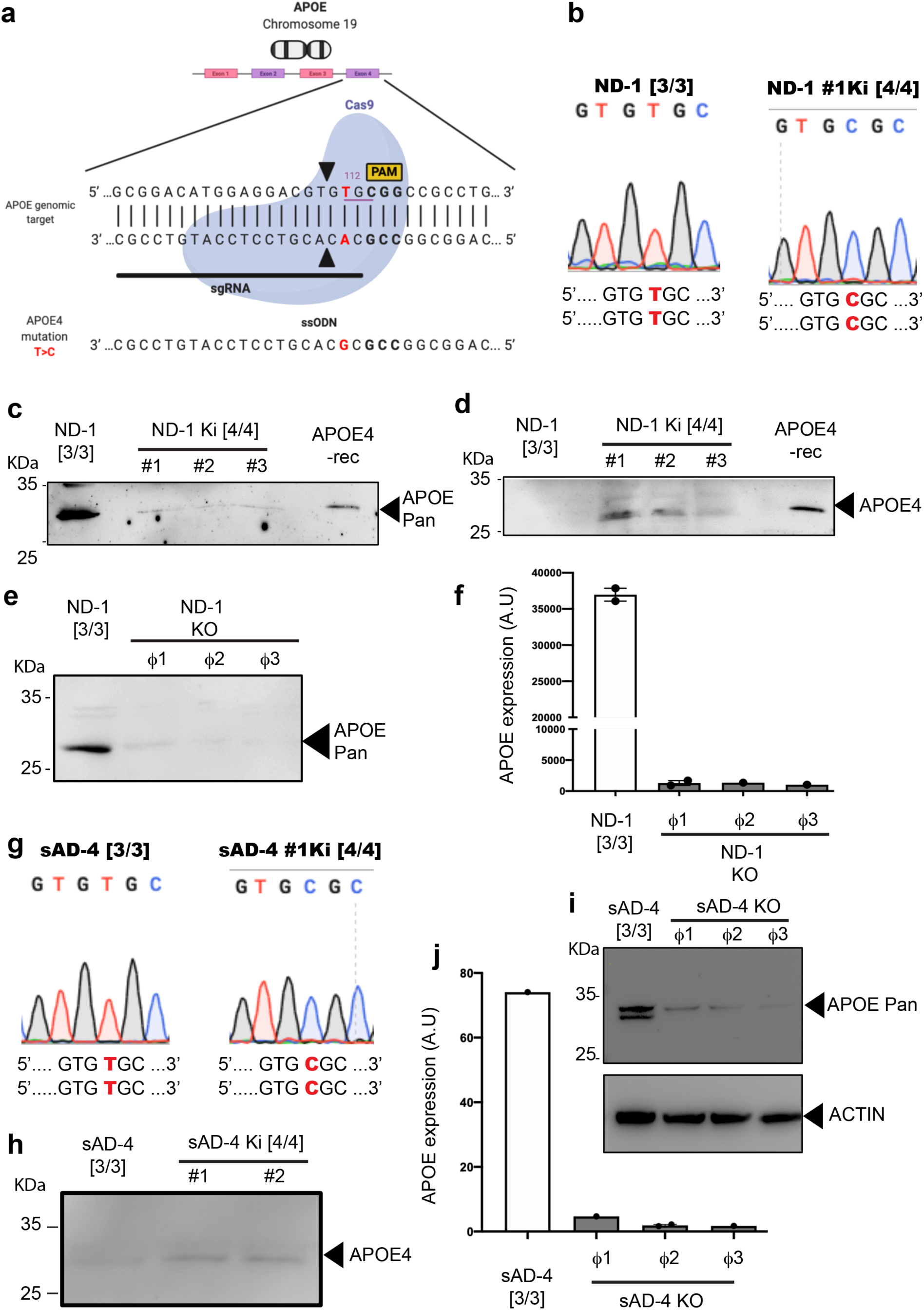
Generation and validation of isogenic control hiPSC lines. **a**) Schematic diagram of the CRISPR-Cas9-based editing strategy used to convert [*APOE3/E3*] hiPSCs, from a non-demented donor (ND-1, b-f) and a sporadic patient diagnosed for AD (sAD-4, g-j), into [*APOE4/E4*]-Knock in (Ki) and *APOE* Knock-Out (KO) hiPSCs. **b**) Traces of Sanger sequencing showing the conversion of a non-demented [*APOE3/E3*] hiPSC line (ND-1 [3/3] (left)) into an isogenic [*APOE4/E4*]-Ki hiPSC clone (right). **c,d**) After Sanger sequencing validation, [*APOE4/E4*]-Ki clones were analysed by Western blot using either an anti-APOE (pan antibody) recognizing the APOE independently of its isoform (**c**) or an antiAPOE4 that recognize the APOE4 isoform specifically (**d**). The APOE4 recombinant protein (APOE4-rec) was used as control. **e**) After Sanger sequencing validation, *APOE*-KO clones (i.e., those displaying major deletions in the targeted sequence) were differentiated into astrocytes and the cell lysates analysed by Western blot using an anti-APOE (pan antibody). **f**) Quantitative analyses (normalized by ACTIN) from Western bloting analyses on *APOE*-KO clones and their control line. **g**) Traces of Sanger sequencing showing the effective conversion of the sAD-4 [3/3] line (left) into an isogenic [*APOE4/E4*]-Ki hiPSC clone (right). **h**) After Sanger sequencing validation, [*APOE4/E4*]-Ki clones and the original sAD-4 [*APOE3/E3*] line were analysed as in **d**). **i,j**) After Sanger sequencing validation (not shown), *APOE*-KO clones (i.e. those displaying major deletions in the targeted sequence) were differentiated into astrocytes, analysed by Western blot (**i**) and further quantified (**j**) using an anti-APOE (pan antibody). In **f** and **j**, A.U. stands for arbitrary units (see methods).

**Extended Data Figure 3.**
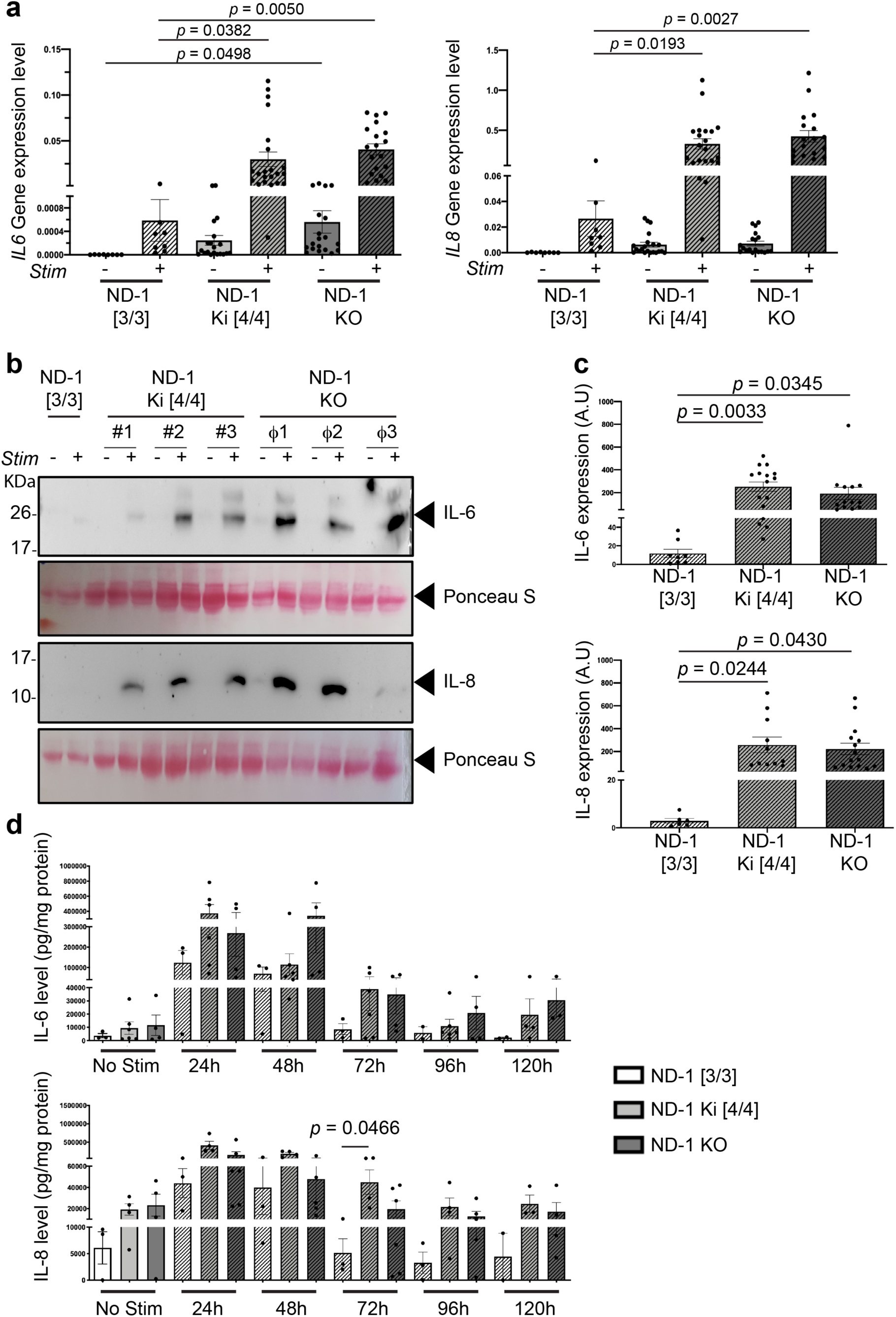
Isogenic APOE4-Ki and APOE-KO clones demonstrate the key role of APOE in the inflammatory response and chronicity from human astrocytes. hiPSCs from a non-demented [*APOE3/E3*] control donor (ND-1 [3/3]) and its respective isogenic [*APOE4/E4*]-Ki (ND-1 Ki [4/4/]) and APOE-KO (ND-1 KO) clones (see also Extended Data Figure 2) were differentiated into astrocyte-like cells prior to challenging with a defined pro-inflammatory cocktail (IL-1β, MCP-1 and TNF-α, referred as Stim). 24 hours after stimulation (+), the cells and the supernatants were collected and processed for analyses of inflammatory mediators by qPCR and Western blotting analyses, respectively. Non-stimulated cultures (−) were also analysed. **a**) Gene expression levels for *IL6* (left) and *IL8* (right) from astrocytes of the indicated genotype. Results are shown as a pool from different clones. **b**) Representative Western blots for IL-6 and IL-8 from [*APOE3/E3*], [*APOE4/E4*]-Ki and *APOE*-KO astrocytes. Ponceau staining was used as loading control for quantitative analyses shown in **c**). **c**) Quantification of IL-6 and IL-8 from Western blotting analyses. Results are shown as a pool from different clones. **d**) Non-demented [*APOE3/E3*] control (ND-1 [3/3]) and its respective isogenic [*APOE4/E4*]-Ki and *APOE*-KO astrocytes were challenged with our proinflammatory cocktail. Using wash out experiments, the daily net production of IL-6 and IL-8 was measured by ELISA every 24 hours over a 120 hours course. Values are from the ND-1[*APOE3/E3*] line and three independent isogenic clones [*APOE4/E4*]-Ki and [*APOE*-KO], with 2 to 5 independent batches of differentiation for each line (**a-c**). For **d**, Values are from one [*APOE3/E3*] line, three isogenic [*APOE4/E4*]-Ki and three *APOE*-KO clones, with 4-9 independent batches of differentiation for each line/clone. Data are presented as mean ±SEM using one-way ANOVA followed by post hoc Dunnett’s test. In c, A.U. stands for arbitrary units (see methods).

**Extended Data Figure 4.**
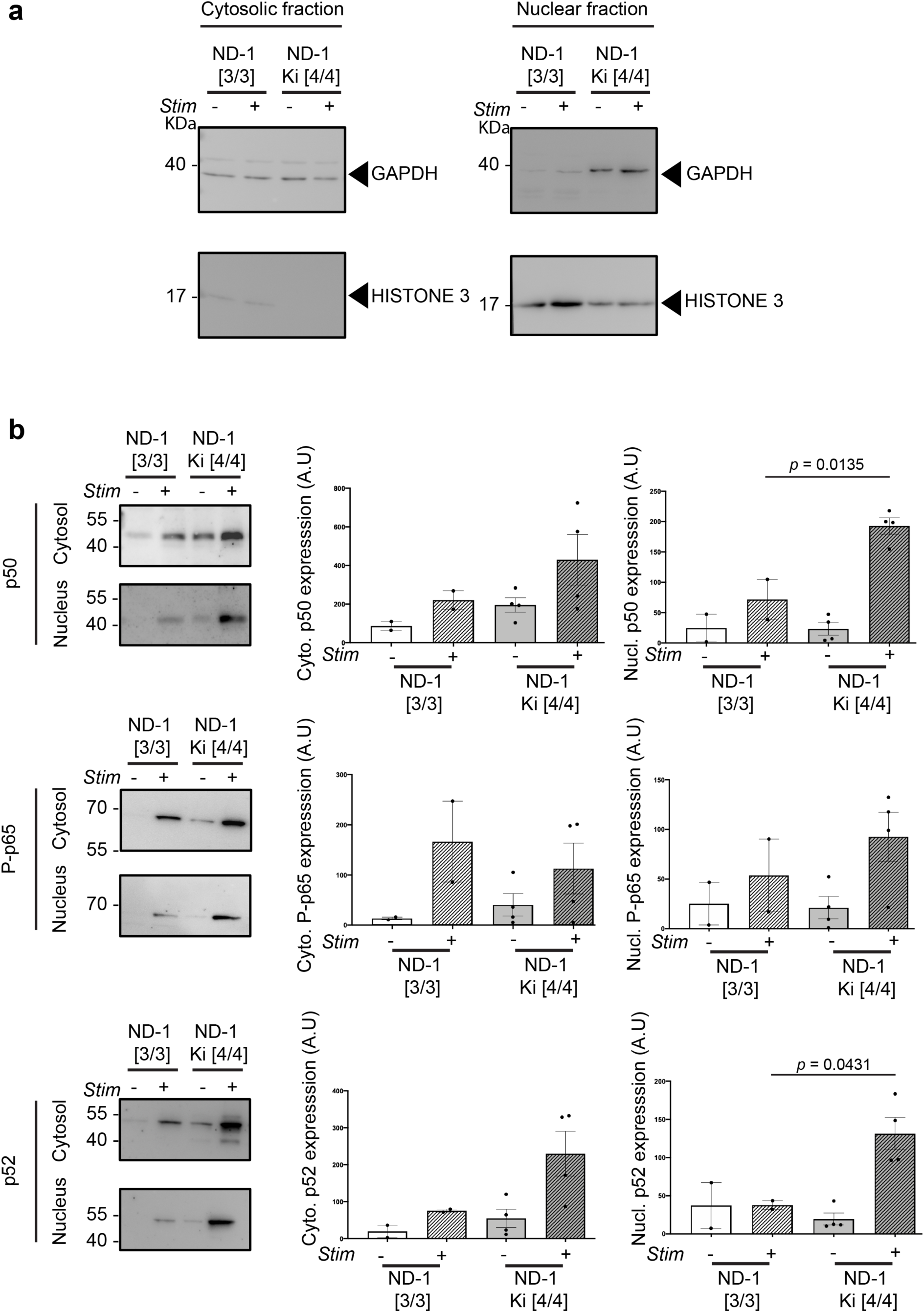
NF-κB activity underlies the APOE4-specific proinflammatory phenotype in astrocytes. Cell lysates from [*APOE3/E3*] (ND-1 [3/3]) and their respective [*APOE4/E4*]-Ki isogenic astrocytes (ND-1 Ki [4/4/]), either in unstimulated or stimulated (Stim) conditions, were prepared for cell fractioning. In **a**) representative Western blots to validate the fractioning (Cytosolic and nuclear fractions). Note that the nuclear Histone 3 was almost exclusively detected in the nuclear fraction, as expected. **b**) From the two fractions, different members of the NF-κB cascade, namely p50 (top), Phosphorylated p65 (P-p65, middle) and p52 (bottom) were analysed by Western blots. Representative immunoblots (left panels) and quantitative analyses (bar charts, right) are shown. Astrocytes from the original ND-1 [*APOE3/E3*] line and two independent isogenic clones [*APOE4/E4*]-Ki were analysed, with 2 independent batches of differentiation for each line. Data are presented as mean ± SEM using unpaired two-tailed Student’s t-Test. In **b**, A.U. stands for arbitrary units (see methods).

**Extended Data Figure 5.**
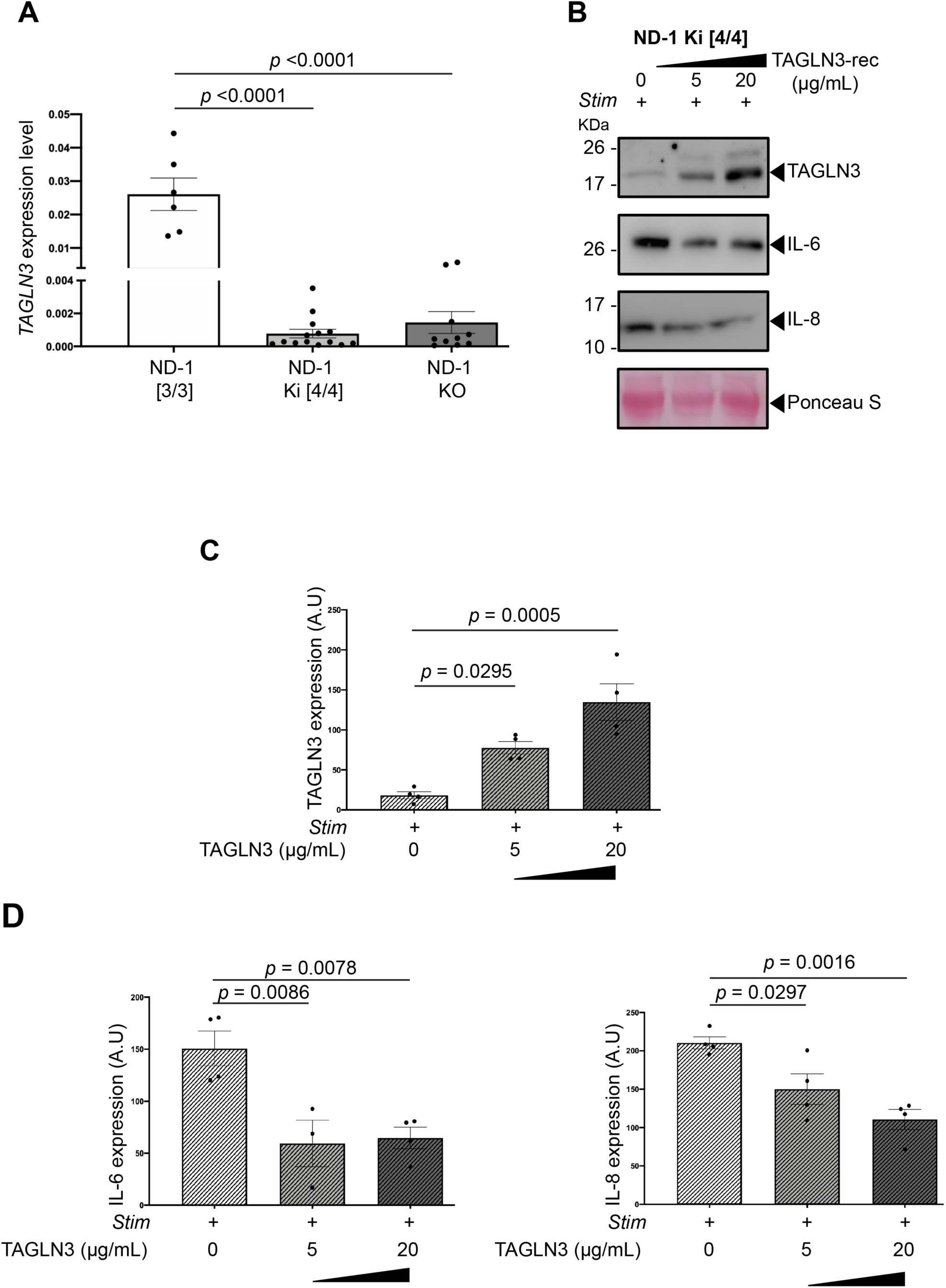
TAGLN3 stands as a new neuroinflammatory modulator in human astrocytes. **a**) Quantification of *TAGLN3* mRNA levels, using qPCR analyses, from [*APOE3/E3*], [*APOE4/E4*]-Ki and [*APOE*-KO] astrocytes. **b-d**) Stimulated [*APOE4/E4*]-Ki astrocytes were treated with 5 or 20 µg/mL of TAGLN3, as recombinant protein (TAGLN3-rec), or with its control vehicle, 24 hours prior to being analysed by Western blot analyses for TAGLN3, IL-6 and IL-8 (**b**). Quantification of Western blot analyses are shown in **c** and **d**. In all different assays, astrocytes from the original ND-1 [*APOE3/E3*] line and three independent isogenic clones [*APOE4/E4*]-Ki and APOE-KO (when applicable) were analysed, with 3 to 7 independent batches of differentiation for each line. Data are presented as mean ±SEM using one-way ANOVA followed by post hoc Dunnett’s test (a, c and d). In **c** and **d**, A.U. stands for arbitrary units (see methods).

**Extended Data Figure 6.**
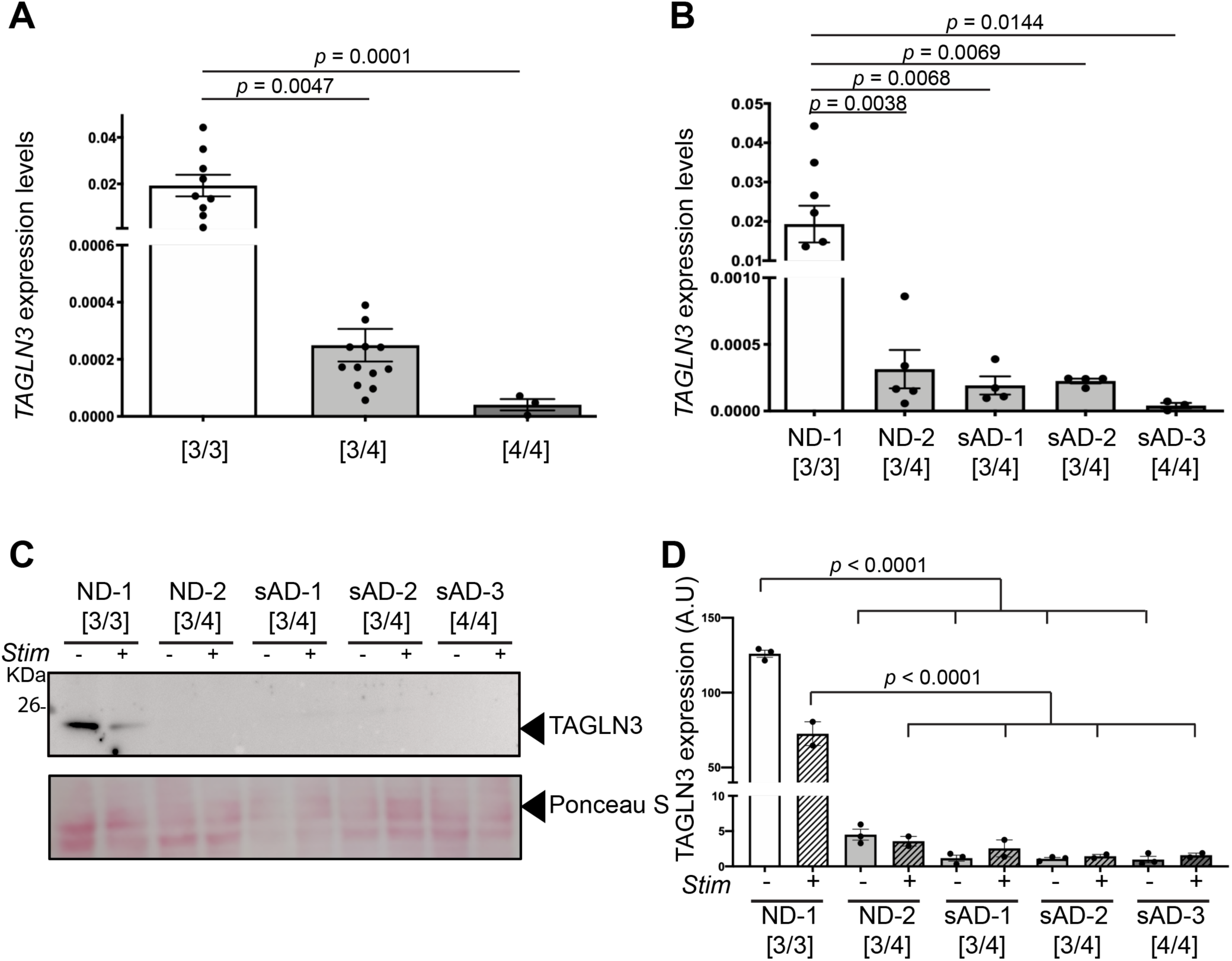
TAGLN3 downregulation in patient-specific astrocytes carrying the *APOE4* allele. **a,b**) *TAGLN3* mRNA expression levels from astrocytes derived from hiPSCs of individuals carrying different APOE polymorphisms [*APOE3/E3*], [*APOE3/E4*] or [*APOE4/E4*]. For each genotype, the *TAGLN3* expression of different hiPSC lines and different batches of differentiation were pooled (**a**) or represented in a patient/donor-specific manner (**b**). **c,d**) Western blotting analyses for TAGLN3 from hiPSC-astrocytes from individuals carrying different APOE polymorphisms [*APOE3/E3*], [*APOE3/E4*] or [*APOE4/E4*]. Non-stimulated (Stim-) and stimulated (Stim+) astrocytes were analysed. Representative Western Blot (**c**) and quantitative analyses (**d**) are shown. For each hiPSC line, 2 to 3 independent batches of differentiation were analysed. Data are presented as mean ± SEM using one-way ANOVA followed by post hoc Dunnett’s test. In **d**, A.U. stands for arbitrary units (see methods).

**Extended Data Figure 7.**
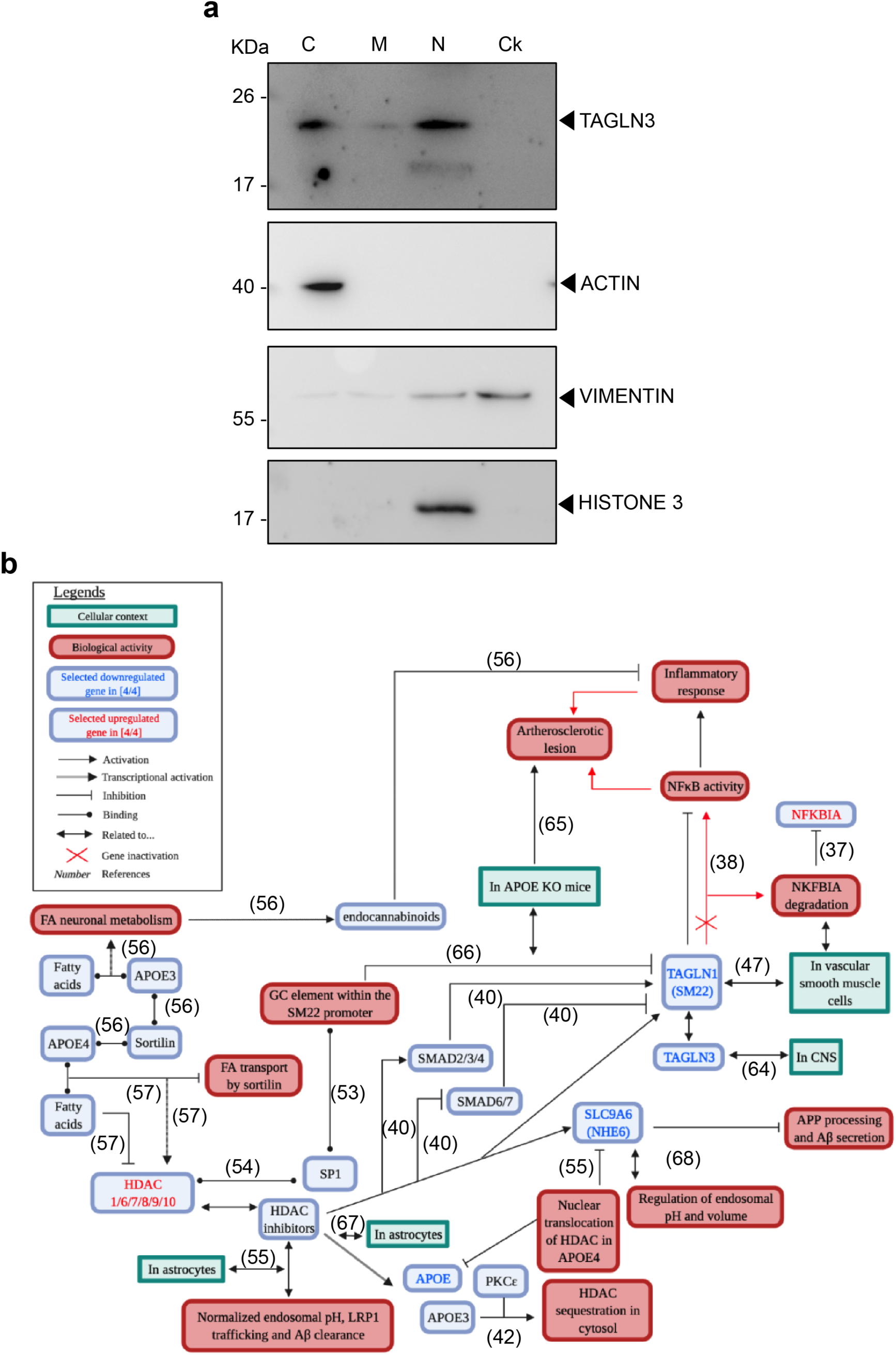
TAGLN3 intracellular distribution and a comprehensive gene interactions map underlying the novel APOE4-TAGLN3 axis. **a**) Representative western Blot revealing the presence of TAGLN3 in in the cytosolic (C) and nuclear (N) fractions of [APOE3/E3] astrocytes. TAGLN3 was barely detected in the membrane (M) fraction and totally absent from the cytoskeleton (Ck) fraction. ACTIN, VIMEMTIN and HISTONE 3 were used as positive control of the C, Ck and N fractions respectively. **b**) Gene interactions map revealing TAGLN3 downregulation as a novel candidate controlling inflammation in an APOE-dependent manner. [APOE3/E3] astrocytes and their respective isogenic [APOE4/E4]-Ki astrocytes, either nonstimulated (No Stim) or stimulated (Stim) with a pro-inflammatory cocktail, were prepared for DNA microarray analysis. Raw data were analysed to identify upregulated and downregulated genes in each experimental condition. Fold changes (FC) cut-off used for analyses were ≥1.5 and ≤0.6 for upregulated and downregulated genes, respectively. Leveraging on those analyses and using PredictSearch® as a powerful text mining software, we established a gene network linking different genes found dysregulated in an APOE-isoform dependent manner to identify possible mechanisms at play in the pro-inflammatory phenotype observed in astrocytes carrying the APOE4 allele. The numbers indicate references (see main and supplementary references).

